# Non-antibiotic drugs break colonization resistance against pathogenic *Gammaproteobacteria*

**DOI:** 10.1101/2023.11.06.564936

**Authors:** Anne Grießhammer, Jacobo de la Cuesta-Zuluaga, Taiyeb Zahir, Patrick Müller, Cordula Gekeler, Hsuan Chang, Katharina Schmitt, Chiara Planker, Erwin Bohn, Taylor H. Nguyen, Kerwyn Casey Huang, Lisa Maier

**Author notes:** authors equally contributed to this work (Anne Grießhammer, Jacobo de la Cuesta-Zuluaga, Taiyeb Zahir).

## Abstract

Non-antibiotic drugs can alter the composition of the gut microbiome, with largely undefined implications for human health. Here we compared the susceptibility of commensal and pathogenic bacteria to non-antibiotic drugs and found that pathogens show higher drug resistance, which could favor their expansion after treatment. We then developed a model system to screen for drug-microbiome interactions that increase the risk of enteropathogenic infections. Approximately 35% of the >50 drugs we tested increased the abundance of *Salmonella* Typhimurium in synthetic and human stool-derived microbial communities. This was due to direct effects of non-antibiotics on individual commensals, altered microbial interactions within communities and the potential of *Salmonella* to exploit different metabolic niches. Non-antibiotic drugs that favored *Salmonella* expansion *in vitro* also promoted other enteric pathogens and increased *Salmonella* loads in gnotobiotic and conventional mice. These findings may inform future strategies to control pathogen proliferation and to assess individual microbiota-drug-pathogen risks for infection.

## Introduction

The human gastrointestinal tract is vulnerable to invasion by non-resident bacteria, including pathogenic members of the class *Gammaproteobacteria* such as non-typhoidal *Salmonella*, *Shigella*, *Vibrio cholerae*, and enteropathogenic or enterotoxigenic *Escherichia coli*^1^. By preventing pathogen colonization and overgrowth of indigenous pathobionts, the gut microbiome provides protection against intestinal infections. This colonization resistance arises from competitive microbe-microbe interactions and the induction of host immune responses^2,3^. Disruption of the microbiome, for instance, during oral antibiotic usage, increases the infection risk^2^. Consistently, post-antibiotic expansion of *Salmonella* Typhimurium (*S*. Tm) has been demonstrated *in vitro*^3,4^, in mouse models^5,6^, and in clinical studies^7,8^.

Non-antibiotic drugs can collaterally damage the human gut microbiome^9^. At physiologically relevant concentrations, hundreds of non-antibiotic drugs can directly inhibit the growth of commensal gut bacteria^10^. As a result, pharmaceuticals from diverse therapeutic classes can alter the composition and function of this microbial community. This includes commonly prescribed antidiabetic^11^, antihypertensive^12^, and antipsychotic drugs^13–16^. Moreover, drug-mediated changes to the microbiome are often dose dependent^17^, can synergize in multi-medicated patients, and can add up with repeated exposures^18–22^. These side effects may cause gastrointestinal symptoms such as diarrhea and gastrointestinal mucosal injury^2,23^.

Population-based metagenomic analyses have shown that intake of several non-antibiotic drugs is associated with higher loads of pathobionts that can cause severe infections^20^. However, it remains unclear whether pathobiont overgrowth results from direct interactions of drugs with the gut microbiome, or from disrupted host immune responses induced by drug use or associated with disease.

Here, we developed a high-throughput *in vitro* assay to test the hypothesis that non-antibiotic medications disrupt the ability of the gut microbiome to resist invaders by interfering with its composition and function. We focused on the growth of *S*. Tm in synthetic microbial communities treated with antibiotics and non-antibiotic drugs, and screened for compounds that alter *S*. Tm expansion after treatment. We identified 18 drugs that promote *S*. Tm growth and 3 drugs that restrict it. Using drug sensitivity assays and synthetic communities of different composition, we showed that alterations to the taxonomic profiles of the community, together with the metabolic characteristics of *S*. Tm, help explain the successful post-drug expansion of the pathogen. We observed similar effects for other pathogenic *Gammaproteobacteria* species, as well as in complex microbial communities derived from multiple human donors. Selected drugs that strongly enhanced the community invasion of the pathogen *in vitro* also disrupted colonization resistance in gnotobiotic mice colonized with the synthetic community and in conventional mice with a complex microbiome.

## Results

### *Gammaproteobacteria* species are more resistant to non-antibiotic drugs than gut commensal bacteria

Using a previously established experimental setup for commensals^10^, we investigated the direct inhibitory effects of approximately 1200 FDA-approved drugs on five pathogenic *Gammaproteobacteria* species: *Salmonella enterica* serovar Typhimurium (*S*. Tm), *Haemophilus parainfluenzae*, *Shigella flexneri, Vibrio cholerae* and *Yersinia pseudotuberculosis*. Overall, these pathogens showed different sensitivity profiles compared to a panel of commensal gut bacteria (Supplementary Figure 1a). While pathogens and gut commensals were inhibited by a similar number of antibiotics (median ± IQR pathogens = 78 ± 4; commensals = 80 ± 16, two-sided t-test t(10.45) = -0.62, adjusted P value = 0.55), pathogens were affected by fewer non-antibiotics compared to commensals (median ± IQR pathogens = 17 ± 7; commensals = 53 ± 37, two-sided t-test t(14.51) = 6.56 adjusted P value < 0.01; Figure 1a, Supplementary Table 1, 20 µM).

**Figure 1:**
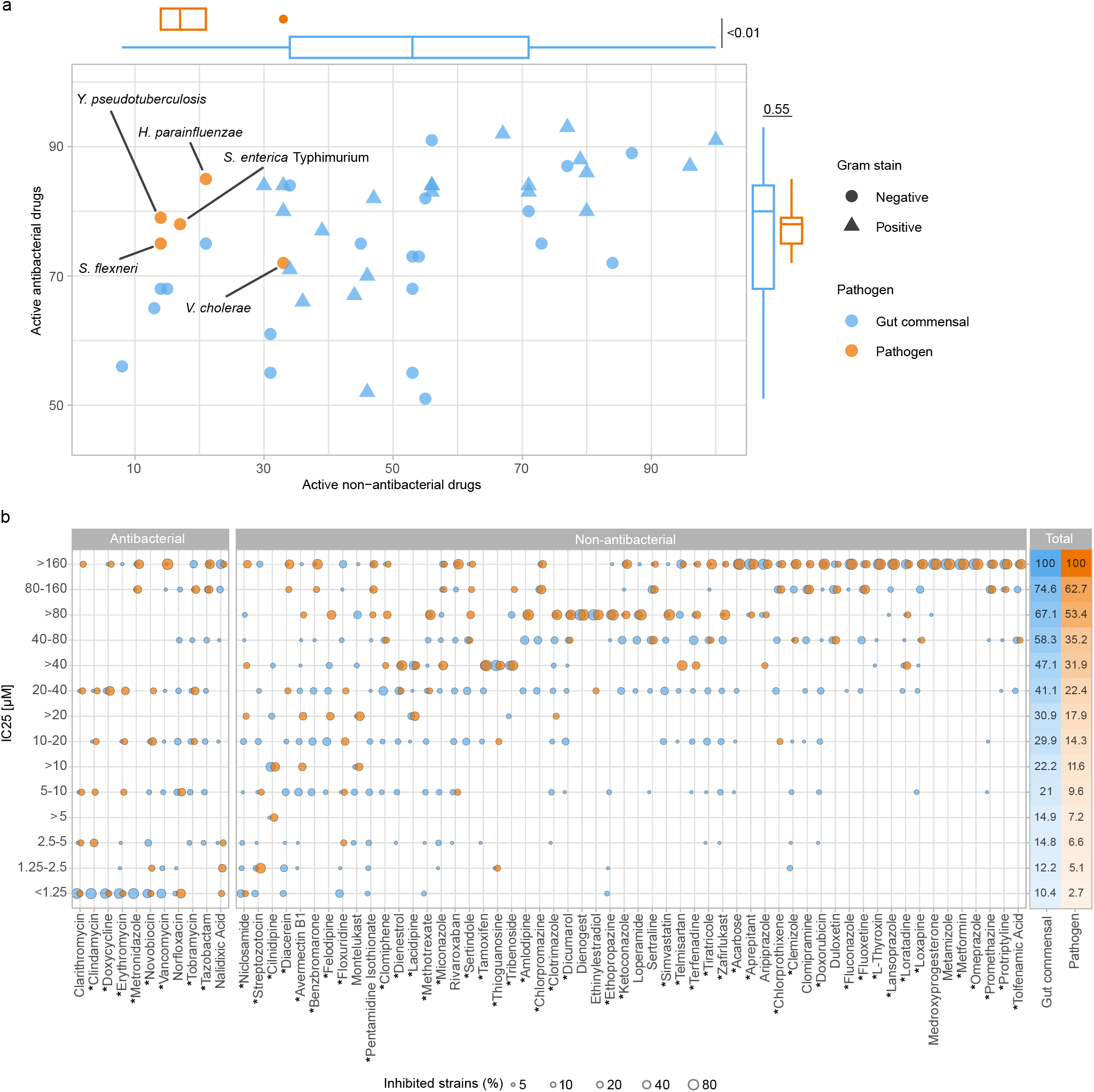
Pathogenic *Gammaproteobacteria* species are more resistant to non-antibiotic drugs than commensal gut bacteria. **a.** Association between the number of active antibacterial and non-antibacterial drugs across 43 gut commensals (blue) and 5 pathogens (orange). Boxplots show the distribution of the number of active antibacterial and non-antibacterial drugs in gut commensals and pathogens. P values from t-tests. **b.** IC_25_ values (25% growth inhibition, Supplementary Table S2) for a panel of 19 gut commensals and 5 pathogenic taxa (*S. enterica* serovar Typhimurium, *S. flexneri*, *V. cholerae*, *Y. pseudotuberculosis* and *Y. enterocolitic*a) across 11 antibacterial and 56 non-antibacterial drugs. Concentrations labeled as greater than (e.g., >20 µM) indicate that the maximum tested did not show any inhibition. Heatmap shows the cumulative proportion of gut commensals and pathogens inhibited at a given concentration. Drugs highlighted with an asterisk (*) were further tested in the *S*. Tm challenge assay.

We further assessed the effect of drugs on gut commensals and *Gammaproteobacteria* pathogens across a wide range of concentrations. For this, we established a panel of 20 bacterial species from the human gut microbiome^24^ and 5 pathogenic *Gammaproteobacteria* species (*Salmonella enterica* serovar Typhimurium, *Shigella flexneri*, *Vibrio cholerae*, *Yersinia pseudotuberculosis* and *Yersinia enterocolitic*a; Supplementary Table 2) and selected 67 antibacterial and non-antibacterial drugs with wide-ranging inhibitory effects across these species in pure culture (Methods, Supplementary Figure 1b). A species was considered to be inhibited by a compound when drug treatment led to a reduction of at least 25% in *in vitro* growth (inhibitory concentration 25; IC_25_). Overall, gut commensals were inhibited at lower concentrations than pathogens by both non-antibiotics and antibiotics: at 20 µM, a physiologically relevant concentration for drugs in the large intestine^10^, 30% of the commensals were inhibited, while only 14% of the pathogens were affected across all drugs (Figure 1b).

These results suggest that differences in susceptibility to pharmaceuticals, and particularly to non-antibiotics, between commensals and pathogens might result in alterations of the gut microbiome that allow pathogenic *Gammaproteobacteria* to thrive, even at low concentrations.

### Microbial community exposure to non-antibiotic drugs modulates *S*. Tm expansion

Next, we assembled a synthetic community comprising the 20 gut commensals tested above. This community, henceforth Com20, is phylogenetically and functionally diverse, spanning 6 bacterial phyla, 11 families, and 17 genera; together, these 20 species encoded up to 61.3% of the metabolic pathways present in healthy human gut microbiomes (Supplementary Figure 2a). Members of the community grew together stably and reproducibly in the gut mimetic medium mGAM^24^, and robustly colonized the gastrointestinal tract of mice for up to 57 days after inoculation of germ-free mice (Figure 2a).

**Figure 2:**
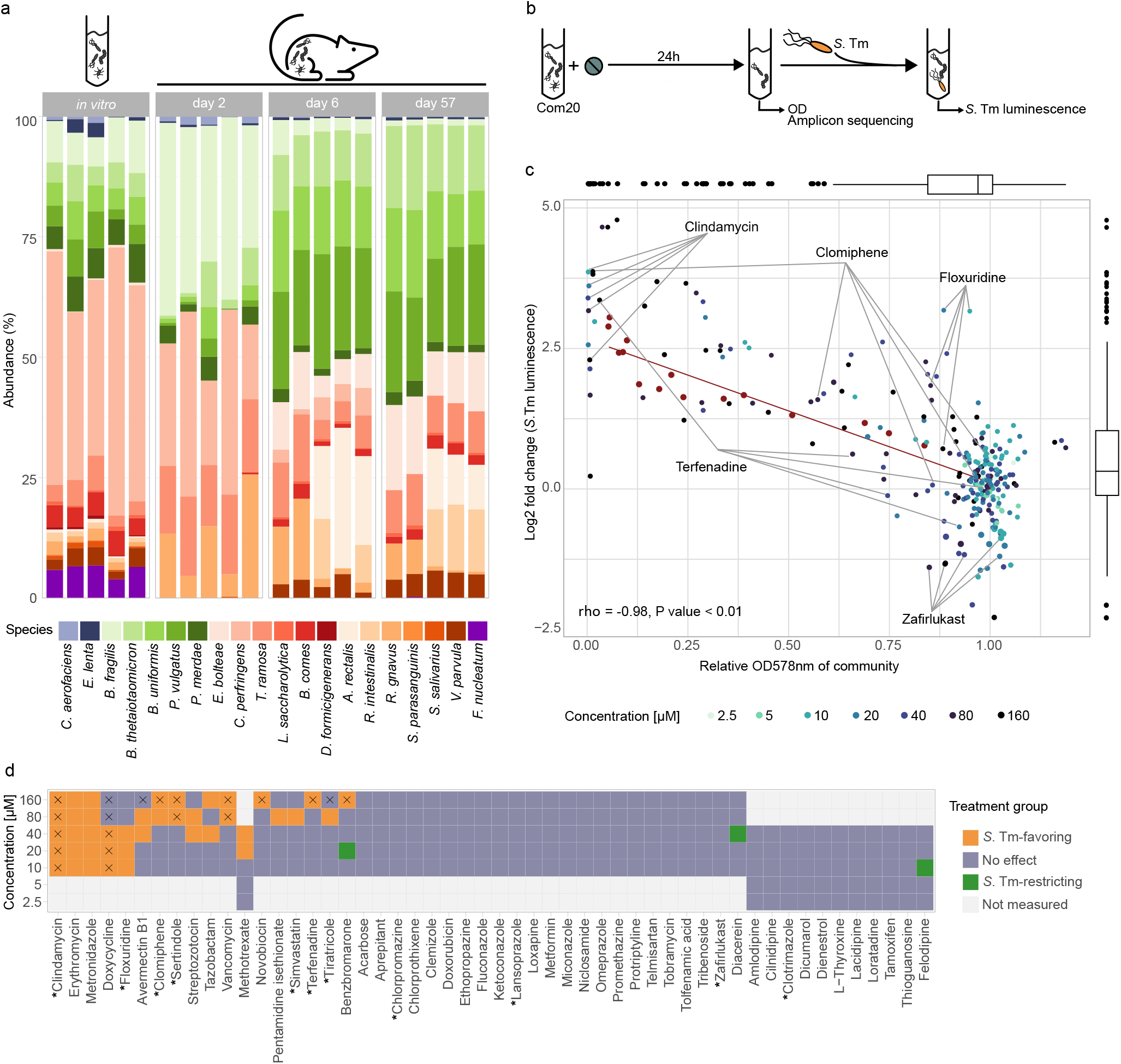
Community exposure to non-antibiotic drugs affects *S*. Tm expansion post-treatment. **a.** Relative abundance of each member of Com20 after 24 h of incubation *in vitro* (1^st^ panel), and in gnotobiotic mice 2, 6, and 57 days after colonization (2^nd^-4^th^ panels). Abundances calculated from 16S rRNA gene sequencing. Each bar represents one biological replicate/fecal sample. **b.** Schematic of the *in vitro S*. Tm challenge assay. Com20 was exposed to drugs across a range of concentrations. After 24 h, optical density at 578 nm (OD_578_) was measured and the drug-treated community was challenged with *S*. Tm in fresh medium. *S*. Tm levels were quantified using *S*. Tm-specific luminescence. **c.** Association between OD_578_ of Com20 and the corresponding *S*. Tm growth in the challenge assay after treatment with 52 drugs. Measurements were normalized to untreated controls. Each point corresponds to the mean of three biological replicates. Red points and regression lines represent the values of untreated, diluted communities. The highlighted drugs were selected for further experiments. **d.** Classification of drugs according to the growth of *S*. Tm in drug-treated Com20 (see Supplementary Figure 2f). Samples marked with an X resulted in a community biomass < 0.2 relative to an untreated community and were excluded from downstream analyses. Drugs labeled with an asterisk (*) were selected for the challenge assay with other pathogenic *Gammaproteobacteria*.

To investigate the effect of drug exposure on the community and *S*. Tm proliferation, we developed a high-throughput challenge assay that mimics the predominance of gut commensal bacteria in the initial stage of community invasion by the pathogen: Com20 was drug-treated for 24 h, after which it was challenged by spiking in *S*. Tm at a small fraction of the biomass of the community (Pathogen to Com20 ratio = 1:500, based on optical density (OD)) in fresh medium. Pathogen growth in Com20 was quantified 4.5 h after challenge using a plasmid-based constitutive luminescence reporter. In absence of any treatment *S*. Tm growth in the synthetic community was lower than in pure culture (*S*. Tm in untreated community: median colony forming units (CFU) 2 x 10^6^ CFU/mL, median relative luminescence units (RLU) 2777 RLU/s. *S*. Tm in pure culture: 2,1 x 10^8^ CFU/mL, 14439 RLU/s) (Supplementary Figure 2b-d). Before pathogen challenge, overall community biomass was measured by OD (Figure 2b). This approach allowed us to differentiate between drugs that altered pathogen growth by reducing overall community biomass from those that altered community composition.

Using this assay, we tested 52 of the 67 drugs previously evaluated in monoculture, at 5 concentrations in triplicate. Drugs inhibiting growth of *S*. Tm were excluded from the challenge assay. Both *S*. Tm luminescence and Com20 OD values were normalized to untreated controls. Measurements were reproducible across replicates (linear regression *R*^2^ = 0.63-0.76 and 0.86-0.92 for luminescence and OD, respectively; Supplementary Figure 2e).

We observed a strong negative correlation between normalized community biomass and normalized *S*. Tm growth: dilution of untreated communities resulted in an increased pathogen growth (Spearman’s rho = -0.98, P value < 0.01; Figure 2c, red dots and line), suggesting that drugs that reduce overall community abundance might open up niches for *S*. Tm. Indeed, antibiotics such as clindamycin favored *S*. Tm growth by strongly decreasing Com20’s biomass, even at the lowest concentration tested.

In contrast, non-antibiotic drugs had a comparatively smaller effect on community biomass, yet resulted in a wide range of different effects on pathogen abundance (Figure 2c, Supplementary Table 3). For instance, certain non-antibiotics, such as the selective estrogen receptor modulator clomiphene and the antihistamine terfenadine, had a concentration-dependent effect on Com20 biomass, resulting in increased *S*. Tm loads in Com20 at higher drug concentrations. Yet other drugs, such as the antimetabolite floxuridine, increased *S*. Tm levels in Com20, despite having only a minor effect on the overall community biomass (Figure 2c, Supplementary Table 3). This observation suggests a second mechanism for drug-mediated pathogen invasion beyond the reduction of Com20 biomass, which is based on changes in its composition. Importantly, changes in the composition of Com20 could not only increase but also decrease *S.* Tm levels, as observed with the calcium channel blocker felodipine and the osteoarthritis drug diacerein (Figure 2d). Similarly, the leukotriene receptor antagonist zafirlukast also tended to reduce *S*. Tm levels in Com20 (Figure 2c, Supplementary Table 3).

To focus on drugs that alter the microbial composition rather than suppressing the complete microbial community, we removed treatments that resulted in a community OD relative to untreated controls < 0.2 from downstream analyses; this led to the removal of 20 treatments corresponding to 10 drugs (Figure 2d). We classified treatments into three groups based on the confidence interval of the mean *S*. Tm growth in treated compared to untreated communities: ‘*S*. Tm-favoring’ (25 concentrations, 11 drugs), ‘*S*. Tm-restricting’ (3 concentrations, 3 drugs) or ‘No effect’ (212 concentrations, 48 drugs) (Figure 2d, Supplementary Figure 2f-g, Supplementary Table 3).

For a subset of treatments, we determined the relative abundance of the commensal species by 16S rRNA gene sequencing post-treatment but before challenge with *S*. Tm (Figure 2b, Supplementary Figure 2h). By normalizing OD to species abundances, we obtained a measure of the contribution of each species to community biomass. Drug treatment resulted in diverse final community compositions depending on the drug used (Supplementary Figure 2i). Community composition post-treatment largely behaved as expected from IC_25_ data: we assessed the ratio of the abundance of each member of the community after treatment to its abundance in a control community and contrasted this to the ratio of growth in monoculture after drug exposure to untreated growth. This allowed us to determine whether the growth of each member in a given community followed the same pattern as in monoculture, be it a reduction or unaffected growth, or whether the microbe was protected or sensitized when part of a microbial community. Across all drugs and all members of Com20, more than 3/4 of drug-microbe interactions followed the IC_25_ results (expected reduced growth = 44.3%; expected unchanged growth = 32.4%). Among the community effects, cross-protection was more frequent than cross-sensitization (protection in community = 19.1%, sensitization in community = 4.2%) (Supplementary Figure 2j), consistent with our previous observations^17^. These results suggest that drug sensitivities in monocultures are largely predictive of the effect of the drug on community biomass, which in turn can be used as an indicator of *S*. Tm expansion in the drug-treated community.

To disentangle the influence of community composition on the growth of *S*. Tm from the effect of the drug on the members of the community, we manually generated four communities whose composition resembled that of drug-treated communities based on 16S rRNA gene sequencing (Supplementary Figure 2k). Specifically, we selected floxuridine (20µM) and zafirlukast (80µM), two drugs that altered *S*. Tm community abundance with minor effect on Com20 biomass, and erythromycin (20µM) and sertindole (80µM) with influence on both Com20 biomass and community composition. We then used these communities to perform a challenge assay. To mimic the change in OD after drug treatment, we performed a serial dilution of the community before spiking in the pathogen. For erythromycin and sertindole, the drug-mimicking conditions did not fully phenocopy treatment. In the case of zafirlukast and floxuridine, drugs that primarily affected *S*. Tm abundance by altering community composition, changing the community composition in the absence of the drug was sufficient to phenocopy the effect on *S*. Tm expansion (Supplementary Figure 2l). These results indicate that changes in the community composition can influence the growth of *S*. Tm in the absence of drug treatment.

Overall, we identified two possible mechanisms by which the effect of drugs on a bacterial community can influence pathogen invasion: first, via a global change in community biomass, and second, via modulation of community composition. For many non-antibiotic drugs, both mechanisms seem to act simultaneously.

### Treatment with non-antibiotic drugs alters the taxonomic profile and metabolic potential of the microbial community

To further assess the association between community composition and *S*. Tm expansion, we identified species whose abundance differed significantly in *S*. Tm-favoring communities compared to untreated Com20 in the set of samples we had amplicon sequencing data. We found that *S.* Tm-favoring communities were enriched in *Collinsella aerofaciens*, *Enterocloster bolteae*, *Dorea formicigenerans*, and *Agathobacter rectalis* (Figure 3a). Conversely, *Phocaeicola vulgatus*, *Bacteroides fragilis*, *Bacteroides uniformis*, and *Streptococcus parasanguinis* were depleted.

**Figure 3:**
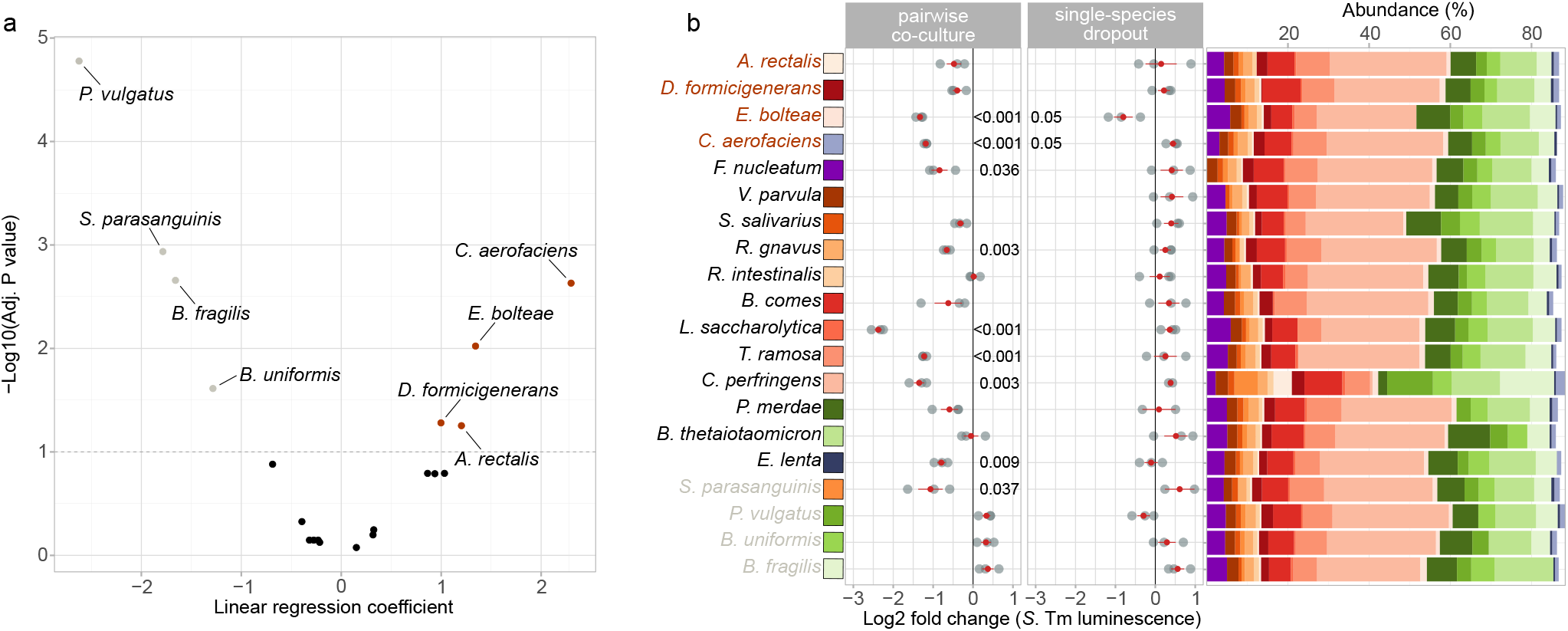
Changes in the community composition and direct commensal-pathogen interactions are connected to *S*. Tm expansion in Com20. **a.** Volcano plot displaying the effect size and adjusted P values of linear regression models of the abundance of members of Com20 in ‘*S*. Tm-favoring’ treatments (n = 33 treatments) compared to untreated controls (n = 45). Dashed line indicates the significance threshold of 0.1. Orange or gray points represent species that significant y increased or decreased, respective y, after ‘*S*. Tm-favoring’ treatments. **b.** *S*. Tm growth in two sets of challenge assays. Left: *S*. Tm luminescence in pairwise co-cultures of *S.* Tm with each member of Com20 (pathogen to commensal ratio = 1:500), compared to *S*. Tm in pure culture. Middle: *S*. Tm luminescence in a 19-member community missing one member of Com20, compared to *S*. Tm in an untreated, 20-member community. Right: the composition of each of the 19-member communities as determined by 16S rRNA amp icon sequencing. Taxa on the *y*-axis are ordered by the regression coefficient from (A). Red points and bars represent the mean ± standard error (SE). Species marked orange were enriched after ‘*S*. Tm-favoring’ treatments, those marked gray were enriched in the controls. Each point corresponds to one of three biological replicates. Raw P values from two-sided t-test; on y values ≤ 0.05 are shown.

We next assessed how interaction of the pathogen with each member of Com20 affected *S*. Tm growth using two complementary approaches: first, a dropout assay, in which we assembled 19-member communities, each missing one of the bacterial species; second, a co-culture assay, in which we co-cultured *S*. Tm with each species (Figure 3b). Community composition remained relatively stable across most dropout assays, with the exception of the community missing *Clostridium perfringens,* which resulted in an increase in *A. rectalis*, *S. parasanguinis* and members of *Bacteroidales.* However, this shift did not translate into changes in *S*. Tm growth (Figure 3b). Moreover, of the 4 species significantly enriched in the *S*. Tm-favoring group, only the removal of *E. bolteae* led to significantly lower pathogen levels, and removal of *C. aerofaciens* even resulted in increased growth. Thus, the growth of *S*. Tm in the drop-out communities did not follow the enrichment patterns observed in the drug-treated communities. The same was true for the pairwise co-culture assays. Based on the enrichment analysis, we expected *E. bolteae* or *C. aerofaciens* to favor *S*. Tm growth, however, both *E. bolteae* or *C. aerofaciens* significantly decreased *S*. Tm growth in co-culture. Together, these observations suggest that *S*. Tm abundance within Com20 is an emergent community property that cannot be fully predicted from the mere presence or absence of single members of the community.

To determine whether differences in the functional potential of the microbial communities correlated with the ability of *S*. Tm to proliferate, we carried out a metagenome prediction of the communities using PICRUSt2. We compared the overall predicted functional capacity of each of the three treatment groups (’*S*. Tm-favoring’, ‘*S*. Tm-restricting’, ‘No effect’) to untreated Com20 and included community OD as a covariate in the model to account for the effect of biomass on *S*. Tm growth. In all cases, we observed significant differences in the overall metagenome potential, indicating that drug treatment results in changes in the functional capacity of the microbiome, independent of its ability to resist pathogens (Supplementary Figure 3a). The largest shifts were observed in the *S*. Tm-favoring group (pairwise PERMANOVA adjusted *R*^2^ = 0.08, 0.05, and 0.02 for *S*. Tm-favoring, No effect, and *S*. Tm-restricting, respectively; adjusted P value < 0.05 in all cases). In addition, compared to untreated Com20 more pathways were significantly different in the *S*. Tm-favoring group than in the *S.* Tm-restricting group (Supplementary Figure 3b, Supplementary Table 4).

In summary, these findings suggest that treatment with non-antibiotic drugs leads to changes in the taxonomic composition and metabolic potential of microbial communities, which are linked to their ability to control pathogen invasion.

### Presence of niche competitor *E. coli* hampers *S*. Tm expansion after drug treatment

Since differences in the metabolic potential of the microbial community were associated with the ability of *S*. Tm to proliferate, we asked whether modifying the niches available to the pathogen influenced its invasiveness. We assessed whether *S.* Tm occupies a distinct metabolic niche from other members of Com20 using predicted genome-scale metabolic models of all species from their genome sequences using CarveMe^25^. We used these models to calculate metabolic competition and complementarity indices with PhyloMint^26^: the metabolic competition index is calculated based on the number of compounds required but not synthesized by both species; the metabolic complementarity index is calculated based on the number of compounds that one species produces that the second species requires but cannot synthesize. We found that *S*. Tm tends to have a lower competition index than most other members of Com20, while having a metabolic complementarity index comparable to that of several members of the community (Figure 4a). Thus, the pathogen can potentially exploit niches that don’t overlap those of the members of the community, while also benefiting from metabolites produced by other bacteria.

**Figure 4:**
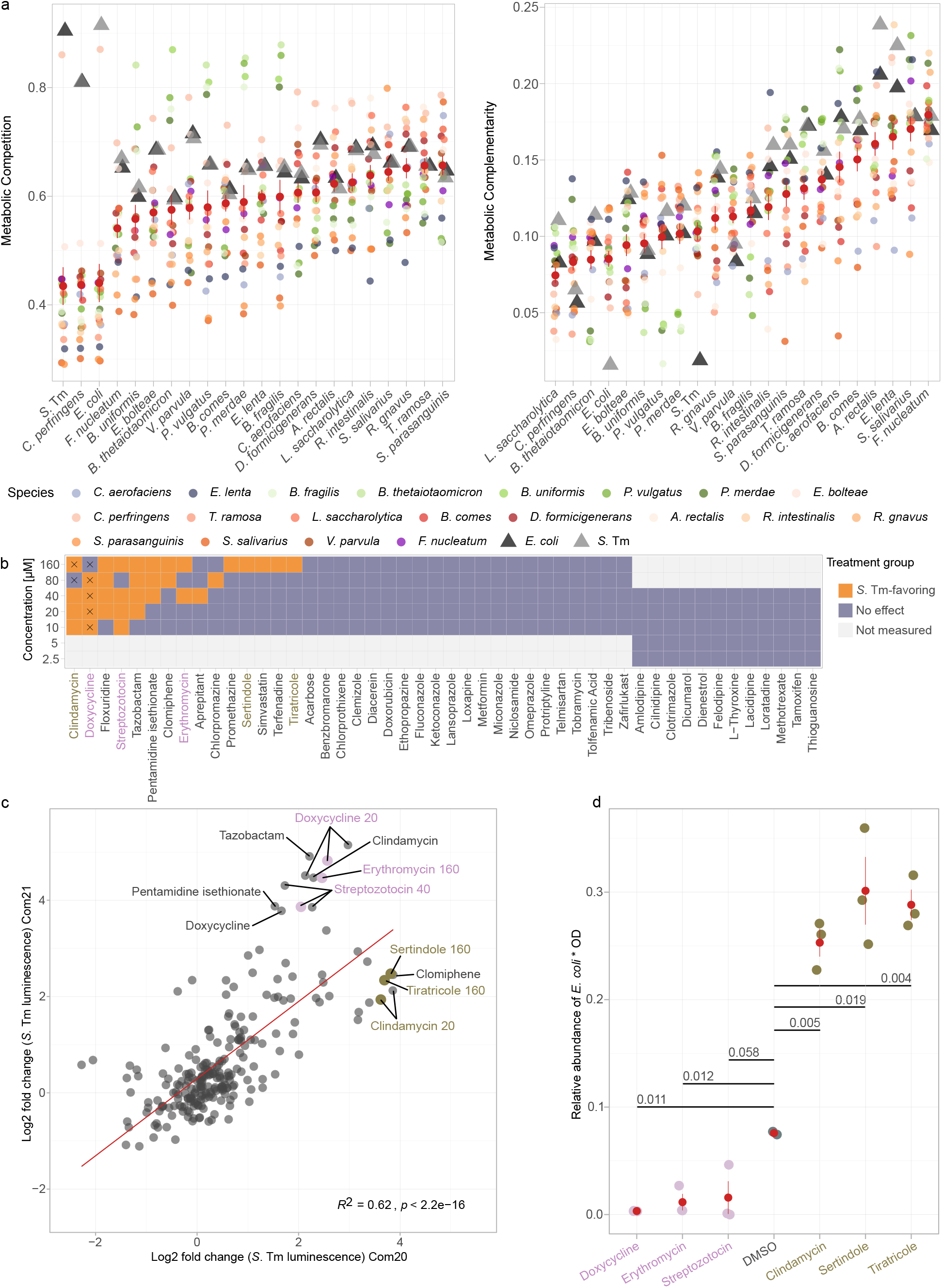
Treatment with drugs that target niche competitor *E. coli* promotes *S*. Tm expansion. **a.** Metabolic competition and complementarity indices of each member of Com21 with each other and with *S*. Tm, calculated from genome-scale metabolic models. Red points and bars represent mean ± SE. Note that the indices are not symmetric. **b.** Classification of drugs according to the growth of *S*. Tm in drug-treated Com21. Conditions that were sequenced are shown in gold and magenta. **c.** Comparison of the post-treatment expansion of *S*. Tm in Com21 versus Com20 across 240 conditions. The red line represents the regression line. Conditions with expansion that differed substantially between the two communities are highlighted (cutoff = 3.5 in log_2_(fold change)), with conditions that were sequenced shown in gold and magenta. **d.** Biomass-scaled relative abundance of *E. coli* in drug-treated Com21. Point colors are based on the groups from (B). Red points and bars represent mean ± SE. Raw P values from unpaired two-sided t-tests are shown.

Based on the above, we reasoned that introducing a bacterium into Com20, which hinders the growth of *S*. Tm by competing for nutrients, would enable us to investigate the role of niche competition in drug-mediated community invasion. *E. coli,* a member of the *Enterobacteriaceae* family with similar metabolic characteristics as *S*. Tm (Figure 4a), is commonly found in the human gut microbiome. Certain *E. coli* strains are known to restrict *Salmonella* species in the intestines of mice and chickens by competing for limiting resources^27,28^. To test whether the presence of *E. coli* in a microbial community inhibits *S*. Tm growth, we added the commensal strain *E. coli* ED1a to Com20, thus generating Com21. The presence of *E. coli* in the microbial community increased the representation of pathways detected in the human microbiome up to 68.6 % (Supplementary Figure 2a). *E. coli-*containing Com21 significantly reduced the growth of *S*. Tm (0.0144 relative *S*. Tm growth to the growth of *S*. Tm alone) compared to Com20 (0.0363 relative *S*. Tm growth to the growth of *S*. Tm alone) (Supplementary Figure 2d). We challenged Com21 with 48 out of 52 drugs evaluated in Com20, at 5 concentrations (Figure 4b); we observed a positive correlation between drug effects on *S*. Tm growth in Com20 and drug effects on *S*. Tm growth in Com21 (Spearman’s rho = 0.62, P value < 0.01; Figure 4c, Supplementary Table 5). However, for some drugs, we observed a different pattern of *S*. Tm growth in Com21 and Com20. In particular, drugs that decreased the abundance of *E. coli* in the community, such as doxycycline, erythromycin, and streptozotocin, led to an increase in *S*. Tm growth in Com21 compared to Com20. Conversely, drugs that increased the abundance of *E. coli*, such as clindamycin, sertindole, and tiratricole, hindered *S*. Tm post-treatment expansion (Figure 4d, Supplementary Figure 4).

These results indicate that drug-induced alterations to the abundance of a niche competitor highly influence the ability of a pathogen to expand within a microbial community. Differences in drug sensitivity of niche competitors will therefore have a substantial impact on invasion outcomes.

### Certain non-antibiotics can also open niches for other *Gammaproteobacteria* pathogens

Other pathogenic members of *Gammaproteobacteria* exhibited similar metabolic requirements (Supplementary Figure 5a) and resistance profiles to non-antibiotics (Figure 1a, Supplementary Figure 1b) as *S.* Tm. Therefore, we asked whether the drugs that influenced post-treatment expansion of *S*. Tm would also affect the growth of other enteropathogens in the absence of closely related competitors. We applied our challenge assay in Com20 to test 12 out of 52 drugs tested before on *S*. Tm, at 5 concentrations on 6 pathogens and pathobionts (Figure 5a, Supplementary Figure 5b). The growth of the other pathogens in drug-treated Com20 was largely consistent with our *S.* Tm results (Figure 5b, Supplementary Table 6). We identified 4 drugs from different classes, namely clindamycin, floxuridine, simvastatin and sertindole, that favored growth across all pathogens tested, including *S*. Tm. Conversely, the effect of certain drugs was pathogen-specific. For example, clotrimazole favored the abundance of all pathogens in Com20 except for *S*. Tm. Treatment with clomiphene led to an increase in pathogen growth except for uropathogenic *E. coli* CFT073 and *V. cholerae* A1552, similar to terfenadine. Chlorpromazine-treated Com20 promoted the levels of *E. coli* CFT073, *S. flexneri* 24570 and *K. pneumoniae* MKP103 but no other pathogens. On the other hand, zafirlukast restricted the growth of *S. flexneri* 24570 *Y. pseudotuberculosis* YPIII, and *Y. enterocolitica* WA-314 but favored the growth of *E. coli* CFT073.

**Figure 5:**
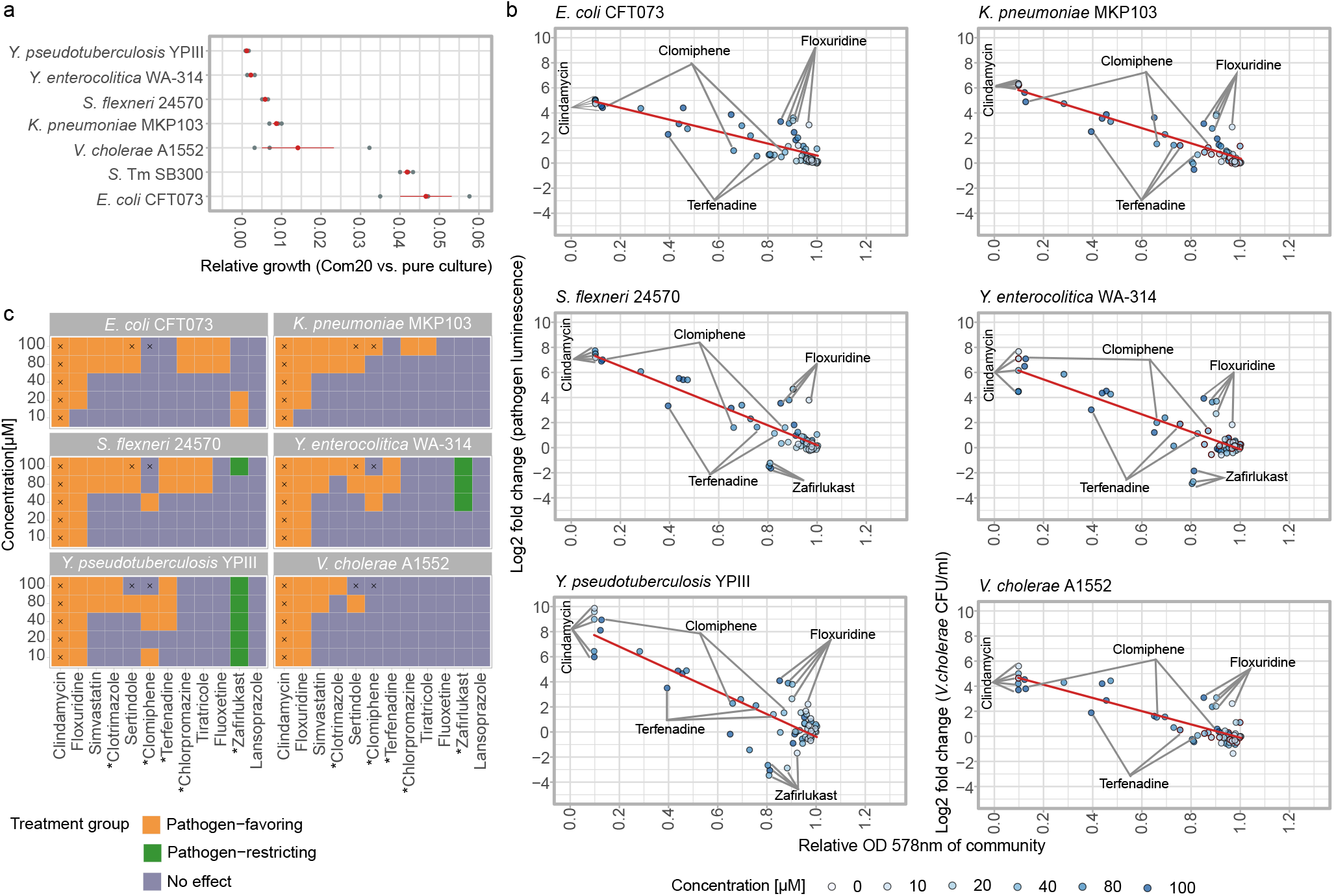
Non-antibiotics that affect *S*. Tm post-treatment expansion also do so for other pathogenic *Gammaproteobacteria*. **a.** Relative growth of various pathogens during co-culture with Com20 compared to pure culture. Pathogen levels were quantified via luminescence, except for *V. cholerae*, which was quantified by selective plating. Each gray point represents one of three biological replicates. Red points and bars represent mean ± SE. **b.** Association between normalized post-treatment OD_578_ and log_2_(fold change) of pathogen luminescence in Com20 for diverse pathogens across 12 drugs at 5 concentrations. *V. cholerae* levels were determined by differential plating instead of luminescence. Each point corresponds to the mean of three biological replicates, point colors indicate drug concentration. Red line represents the regression line. **c.** Classification of drugs according to the growth of each pathogen in Com20. Drugs highlighted with an asterisk (*) were later tested *in vivo*.

These findings indicate that some community-disrupting effects can be generalized to other pathogenic *Gammaproteobacteria* (Figure 5c). However, certain drug-induced changes modulate the growth of specific pathogens, highlighting the importance of considering pathogen-specific aspects of colonization resistance.

### *S.* Tm expansion in human-stool-derived communities after non-antibiotic treatment recapitulates *in vitro* assays

Our synthetic communities are simplified models. As such, they are advantageous for studies *in vitro* and in gnotobiotic mice, however compared to the human gut microbiome, they have reduced species diversity, lack intraspecies variation and do not recapitulate individual differences. Thus, we sought to test whether drug treatment of complex and microbially diverse communities resulted in similar expansion of *S*. Tm. For this, we derived stable microbial communities^3,29^ from fecal samples from 8 healthy adults, which we used for *in vitro* assays. We found that the sensitivity of the stool-derived communities to diverse drugs varied across donors (Supplementary Figure 6), suggesting inter-individual differences in how these communities respond to drug exposure. Next, for 8 drugs at various concentrations, we evaluated the ability of *S.* Tm to grow in these communities after treatment. *S.* Tm growth in Com20 and Com21 was positively correlated with growth in the stool-derived communities (Figure 6a, Supplementary Table S7). Notably, the mean correlation across donors was higher for Com21 than Com20 (Spearman’s rho = 0.57 and 0.74 for Com20 and Com21, respectively; Figure 6b), and the number of significant donor-community correlations was higher with Com21 (8 and 5 for Com21 and Com20, respectively; Figure 6b). The increased similarity between the response of Com21 and stool-derived communities compared to that of Com20 might be explained by the presence of the genus *Escherichia*, which strongly increased the representation of metabolic pathways present in the human gut microbiome (Supplementary Figure 2a). Similar to the differences in drug sensitivity between the different communities, *S*. Tm levels after treatment with non-antibiotics varied between individuals (Figure 6). This suggests that the effect of drugs on S. Tm invasion can be further modulated by individual community properties.

**Figure 6:**
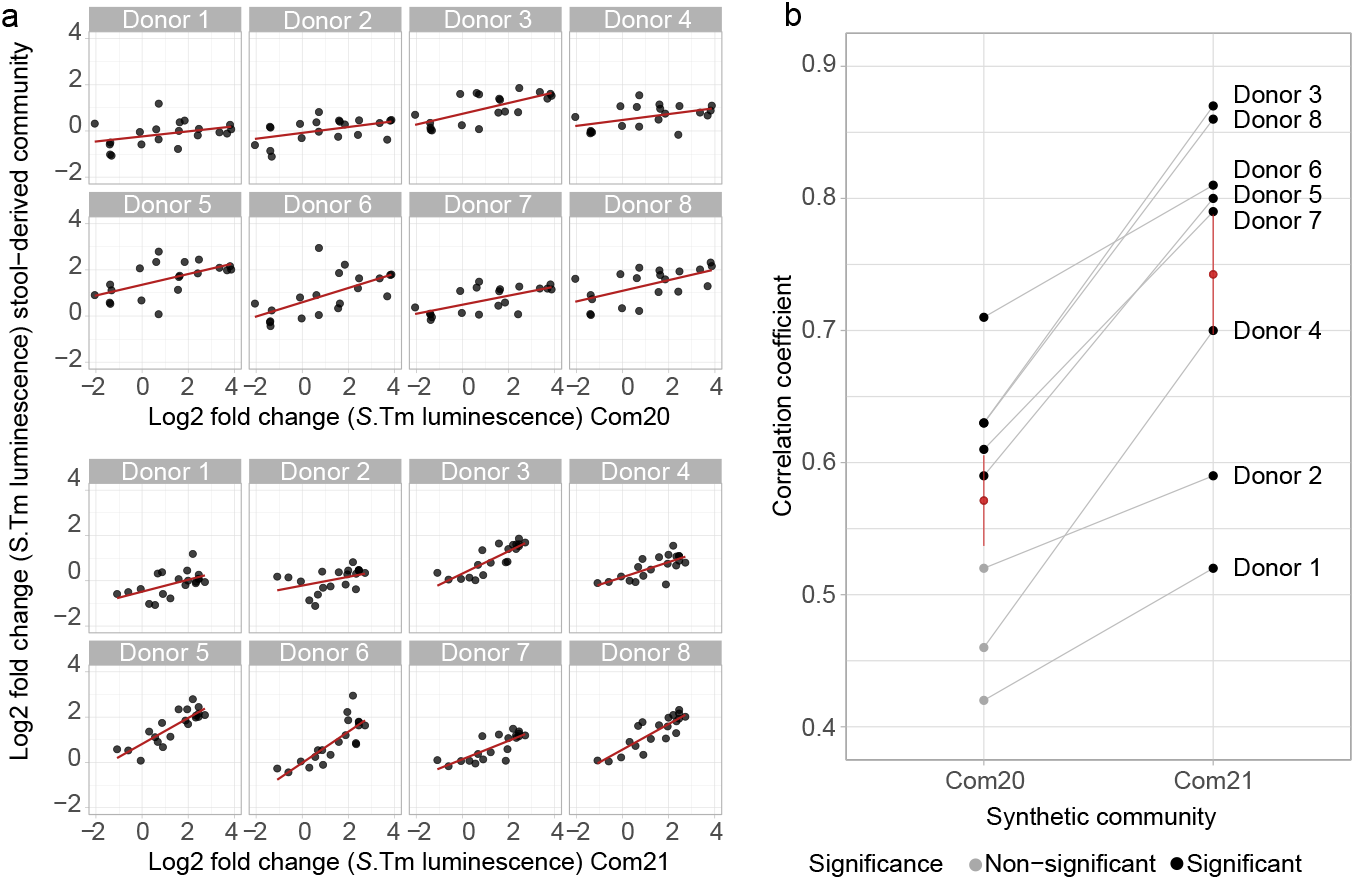
Drug-induced expansion of *S*. Tm is reproducible in complex communities derived from human feces. **a**. Association between *S*. Tm luminescence in drug-treated Com20 (top) and Com21 (bottom) with pathogen growth in drug-treated stool-derived microbial communities, relative to untreated microbial communities. Regression lines shown in red. Each point corresponds to a drug/concentration combination (n = 20 from s drugs). **b**. Spearman’s correlation coefficient of *S*. Tm growth between individual stool-derived communities and Com20 or Com21 across multiple drugs from (A). Black points represent associations with adjusted P value <0.1. Red points and bars represent mean ± SE across s human-derived communities.

Collectively, these findings support the notion that non-antibiotic drugs can influence the levels of *S*. Tm and other pathogenic *Gammaproteobacteria* species in the human gut microbiome, and that the extent of the shifts induced by treatment with non-antibiotics might vary based on the individual microbiome composition. In addition, these results further validate our synthetic communities as models for studying the effects of these compounds on the human gut microbiome.

### Non-antibiotic drugs disrupt colonization resistance against *S.* Tm in mice

Finally, we assessed whether the modulation of *S.* Tm growth by non-antibiotics observed *in vitro* would translate into a disruption of colonization resistance *in vivo* using two animal models (Figure 7a): first, gnotobiotic mice colonized with Com20 to test the transferability of our *in vitro* work and second, conventional specific pathogen free (SPF) mice to test a complex mouse microbiome (on average, 199 amplicon sequence variants (ASV) before treatment).

**Figure 7:**
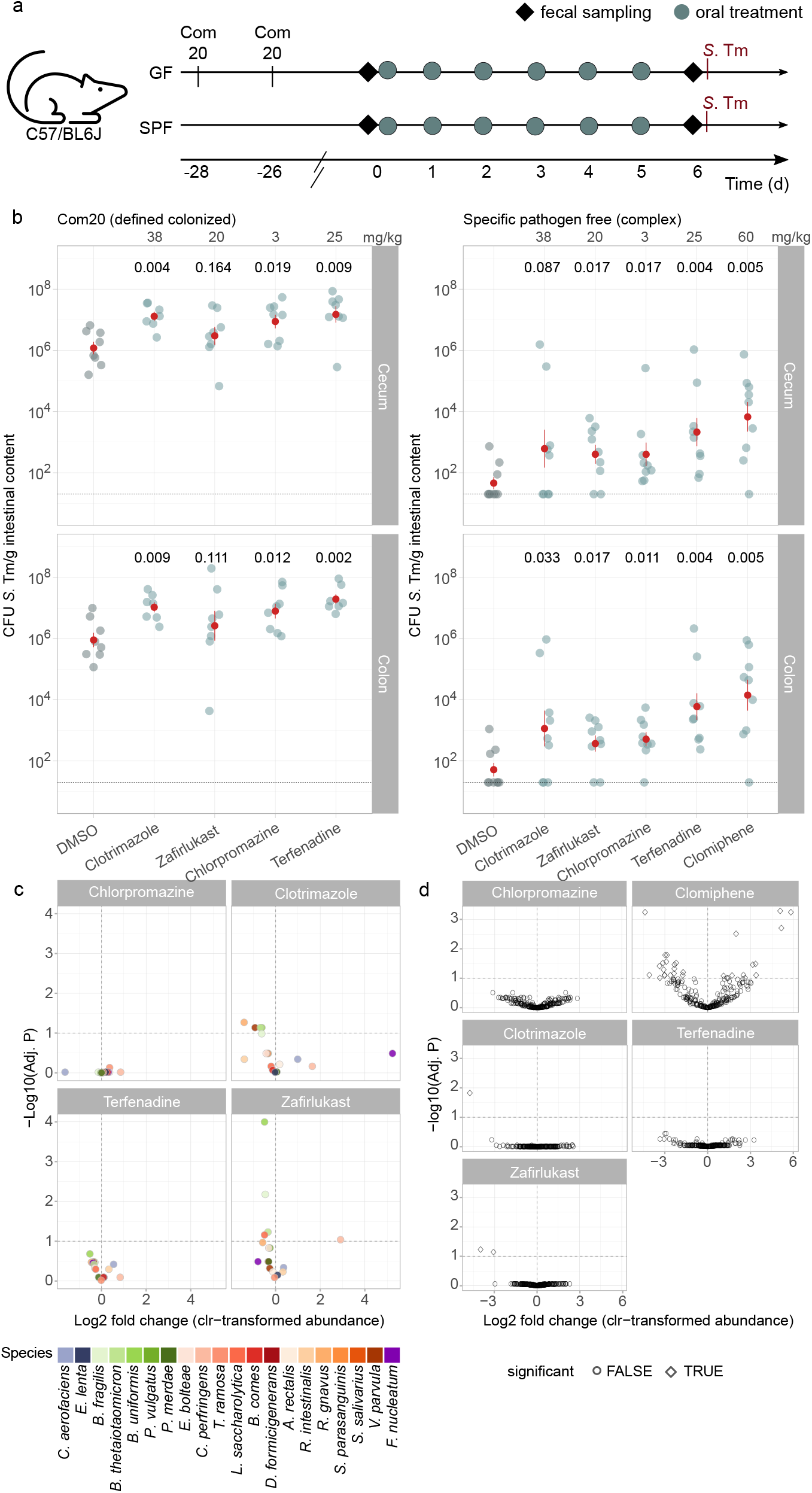
Non-antibiotics from diverse therapeutic classes disrupt colonization resistance in mice. **a**. Schematic of mouse treatment and sampling. **b***. S*. Tm load in cecum and colon on day 1 post challenge after treatment with four drugs for 7 days at the doses indicated (top). Red points show the mean values of 7-9 biological replicates, red lines represent mean ± SE. Adjusted P values from one-tailed Wilcoxon-test with BH correction of the comparison of drug-to vehicle (DMSO)-treated mice. **c.** Volcano plot of the effect size and adjusted P values of linear models of the abundance of Com20 members in gnotobiotic mice 6 days after drug treatment compared to untreated controls, adjusting for species abundance on day 0. Dashed line indicates an adjusted P value significance threshold of 0.1. **d**. Volcano plot of the effect size and adjusted P values of the abundance of amp icon sequence variants (ASVs) in SPF mice 6 days after drug treatment compared to untreated controls, adjusting for ASV abundance on day 0. Dashed line indicates an adjusted P value significance threshold of 0.1. **e**. Point shape indicates whether an ASV was significant y different (TRUE) from controls.

We first validated these models by assessing *S*. Tm growth after a single dose of 800 mg/kg streptomycin, which was expected to disrupt colonization resistance^5^. Twenty-four hours post challenge, pathogen levels reached 10^8^-10^9^ colony forming units (CFU) per gram of feces in both models (Supplementary Figure 7a), confirming their suitability for investigating drug effects on *S*. Tm colonization.

Next, we selected 5 drugs with distinct effects *in vitro* (Figure 2c and 2d): clotrimazole (No effect on *S*. Tm but on most other pathogens), chlorpromazine (No effect on *S*. Tm expansion in Com20 but in Com21, pathogen-favoring on most other pathogens), zafirlukast (No effect on *S*. Tm but restricting other pathogens), and terfenadine and clomiphene (favoring most pathogens, including *S*. Tm). We administered these drugs to mice at concentrations equivalent to the human dose for chronic treatment (3-60 mg/kg, Figure 7a). In Com20-colonized mice, most tested drugs resulted in significantly higher *S*. Tm levels compared to the DMSO control, with the exception of zafirlukast, which was expected given our *in vitro* results (Figure 7b). Interestingly, clotrimazole and chlorpromazine, which only showed a significant effect in the *in vitro* assay for pathogens other than *S*. Tm, resulted in higher *S*. Tm levels *in vivo*, suggesting that the contribution of non-antibiotics to the disruption of colonization resistance may be underestimated in our *in vitro* data. In SPF mice, all drugs, including zafirlukast, resulted in significantly higher *S*. Tm levels, indicating that the effect of zafirlukast might be dictated by the community and/or host context (Figure 7b).

In either mice model, the composition of the microbiota after drug treatment compared with untreated controls and adjusting for baseline taxa abundances did not undergo extensive changes. In gnotobiotic mice, only treatment with clotrimazole and zafirlukast resulted in significant changes in the abundance of 4 and 5 species, respectively (Figure 7c, Supplementary Figure 7b, Supplementary Table S8). Similarly, 31, 2 and 1 amplicon sequence variants (ASVs) were significantly different in SPF mice treated with clomiphene, clotrimazole and zafirlukast, respectively (Figure 7d, Supplementary Table S9). This implies that communities are more protected from drugs *in vivo*, possibly by emergent behaviors in the host context and reduced effective drug concentrations in the colon. Regardless of statistical significance, the change in abundance of certain taxonomic groups was consistent across treatments. In gnotobiotic mice, drug treatment tended to increase the abundance of *E. lenta*, *C. aerofaciens*, and *C. perfringens* (Supplementary Table S8). In SPF mice, the proportion of ASVs from the orders *Lachnospirales* and *Oscillospirales* that showed changes in abundance was comparable, however, more ASVs from the order *Bacteroidales* tended to increase than decrease in abundance in treated mice (Supplementary Table S9).

We did not observe any significant changes in *E. coli* counts in SPF mice after drug treatment (Supplementary Figure 7c), suggesting that despite the presence of a close competitor (Figure 4a), *S*. Tm has a growth advantage in drug-disturbed microbial communities. In none of the mouse models did higher *S*. Tm loads lead to host symptoms, signs of intestinal inflammation or systemic infection 24 h after *S*. Tm challenge (Supplementary Figure 7d). This is consistent with previous reports indicating that *S*. Tm loads in the range of 10^8^ CFU/ g feces are required to trigger inflammation^5^.

Overall, our results confirm our *in vitro* findings by showing that non-antibiotic drugs from different therapeutic classes abrogate colonization resistance against *S*. Tm in mice with defined and complex microbiotas. For certain drugs, the interference with colonization resistance depended on the specific microbiome composition, as shown for zafirlukast.

## Discussion

In this study, we systematically investigated how drug-induced disruption of microbial communities affected pathogen invasion *in vitro* and *in vivo*. Starting from a set of more than 1200 compounds, our approach led to the identification of non-antibiotic drugs that increased *S*. Tm pathogen load in gnotobiotic and conventional mice. These drugs included compounds from a wide range of therapeutic classes, including anti-asthmatic, antipsychotic, selective estrogen receptor modulators, antifungal and antihistaminic drugs. However, we also discovered drugs, such as zafirlukast, which hindered the expansion of multiple pathogenic *Gammaproteobacteria* species *in vitro*. Overall, we found more compounds that favored *S*. Tm expansion rather than restricted it, consistent with our observation that *Gammaproteobacteria* were more resistant to drugs.

To the best of our knowledge, the present study is the first to demonstrate that non-antibiotic drugs can disrupt colonization resistance, increasing the burden of pathobionts and pathogens. This is consistent with a population-level metagenomic cohort study in which an association between non-antibiotic drug consumption and higher pathobiont load was reported^20^. This implies a previously underestimated risk of infection, with potentially severe consequences in vulnerable populations such as immunocompromised individuals or those taking multiple or chronic non-antibiotic medications.

Our approach is based on 20/21-member model communities, which are ideally suited to dissect intricate species-species and drug-species interactions within the human gut microbiome. The members of these communities are prevalent and abundant gut bacterial species and have been thoroughly characterized for the direct interaction across 1200 marketed drugs^10,30^. While only encompassing a small number of species, both communities encoded roughly two thirds of the metabolic pathways detectable in the human gut microbiome by state-of-the-art metagenome profilers^31,32^. Several properties make these communities an attractive model to study colonization resistance. First, in addition to its utility for *in vitro* analyses, this community stably colonized the mouse gut. Second, the overall community composition was similar both *in vitro* and *in vivo*, with the dominant members belonging to the phyla *Bacteroidota* and *Bacillota*, the main phyla in the human gut microbiota. Third, without drug treatment, Com20/21 hindered the growth of *Gammaproteobacteria* pathogens, recapitulating key features of colonization resistance. Fourth, the outcome of the *S*. Tm challenge assays using Com20/21 positively correlated with the response observed in human-stool-derived communities. Therefore, although it is a reductionist model, it recapitulates many of the characteristics of a natural microbiome. Our overall strategy makes it possible to disentangle the contribution of individual microbes or microbial consortia from that of the host in preventing pathogen expansion after drug treatment. All the above make our approach a powerful tool for the study of mechanistic aspects of colonization resistance and provide a complement to standard preclinical animal models and cohort studies.

The present study represents a conservative estimate of the number of non-antibiotics with the potential to increase the pathogen load. This is because the *in vitro* assay is unable to measure host-mediated aspects of colonization resistance, such as immunological (e.g., antimicrobial peptides) and physical barriers (e.g., mucus, spatial variation along the intestine), a shortcoming of our approach. Consequently, we may have overlooked non-antibiotic drugs that promote post-treatment pathogen growth due to microbiome-independent factors. One such group are proton-pump inhibitors, which strongly inhibit gastric acid production and are considered a risk factor for antibiotic-induced diarrhea^33,34^.

Using the model communities, we found that non-antibiotic drugs can affect both the composition as well as the overall abundance of microbes, and that both factors are linked to the ability of multiple *Gammaproteobacteria* pathogens to proliferate. According to the nutrient-niche concept, the success of a pathogen in colonizing the intestine depends on its ability to identify a suitable niche and outcompete other members of the ecosystem by efficiently consuming limiting substrates^35,36^. After antibiotic treatment, large shifts in microbiome composition and function lead to an increased release of monosaccharides, facilitating *S*. Tm expansion^23,37,38^ and resulting in the induction of self-promoting host inflammation^23,37–39^. Our work expands upon this concept by incorporating differences in drug sensitivities into the ecological considerations. While antibiotics can affect a very wide range of species, non-antibiotics tend to target specific commensal bacteria. This precise inhibition can have critical consequences for community composition and/or function and might explain why pathogen load increases even after non-antibiotic treatment, although the overall collateral damage is smaller compared to antibiotics.

As a metabolic generalist, *Salmonella* can easily adjust to diverse post-drug nutrient landscapes. However, this adaptability is also a property of direct niche competitors such as commensal *E. coli*^28^, which is similarly resistant to non-antibiotics. Thus, minor differences in drug sensitivities between resident microbes and invading pathogens, combined with their differential ability to utilize limited substrates, will likely influence colonization and, ultimately, infection outcomes. Importantly, the variation in sensitivity to non-antibiotics we observed in stool-derived communities *in vitro* and the variation of these communities to hinder *S*. Tm proliferation suggests that, at a fine scale, inter-individual differences in microbiome compositions influence colonization outcomes.

While it is known that the gut microbiome confers colonization resistance against enteric pathogens^40^, the exact role played by specific commensals is not clear and reports are contradictory. It has been reported that *B. thetaiotaomicron* can exacerbate the colonization by enteric pathogens in mice^41^. Conversely, others have shown that *Bacteroides* species can block *Salmonella* colonization via production of the short-chain fatty acid (SCFA) propionate^42^. The latter finding is consistent with our observations: *B. uniformis*, *B. fragilis*, and *P. vulgatus* were depleted in communities that favored *S*. Tm growth *in vitro* after drug treatment and the abundance of these species was lower on average in drug-treated gnotobiotic mice compared to untreated controls. In contrast, our results with SPF mice showed that members of the order *Bacteroidales* were enriched in drug-treated mice compared to untreated controls, while the orders *Oscillospirales* and *Lachnospirales* exhibited lower abundances. These latter two orders are diverse clades whose members are fiber degraders and SCFA-producers, commonly found in the mammalian gut microbiome^43^, and are largely associated with positive health outcomes in humans^44^. One plausible reason for the apparent inconsistency in *Bacteroidales* outcomes in our *in vivo* and *in vitro* models may be that control of pathogen growth is linked to the levels of short-chain fatty acids, rather than the abundance of the specific taxa responsible for their production.

In summary, the present work emphasizes the risks posed by non-antibiotic drugs in disrupting the microbiome’s ability to protect against pathogen colonization and increasing the likelihood of infections. Our approach provides a basis for understanding the mechanisms of non-antibiotic-mediated expansion of pathogenic bacteria, which is critical for the development of strategies to reduce pathogen burden in vulnerable populations. Future studies examining the effects of non-antibiotic drugs across a wide range of microbiome compositions, drug dosages, and treatment regimens will be pivotal in the development of strategies to predict, mitigate, and minimize microbiome-mediated side effects of these medications.

## Acknowledgments

The authors thank all members of the Maier lab and Athanasios Typas (EMBL Heidelberg) for fruitful discussions and comments on the manuscript, the Foster lab (Oxford, UK) for plasmid pIJ11282 ilux, Libera Lo Presti for proof-reading, Lena Michaelis and Johann Zent for support of our animal work, the Gnotobiotic Research Center Tübingen (GRCT), and the NGS Competence Center Tübingen (NCCT). L.M. acknowledges funding from the DFG (Cluster of Excellence CMFI EXC 2124, Emmy Noether Programme MA 8164/1-1). J.dlC.Z. and L.M. received support from the BMBF-funded de.NBI Cloud within the German Network for Bioinformatics Infrastructure (031A532B, 031A533A, 031A533B, 031A534A, 031A535A, 031A537A, 031A537B, 031A537C, 031A537D, 031A538A). K.C.H. acknowledges funding from NIH RM1 GM135102 and R01 AI147023, and NSF grants EF-2125383 and IOS-2032985. K.C.H. is a Chan Zuckerberg Biohub Investigator. T.H.N. is supported by the NSF Graduate Research Fellowship.

## Authors contributions

Conceptualization: L. M.; Methodology: A. G., T. Z., P. M., J.d.l.C.Z and L. M.; Formal analysis: J.d.l.C.Z, A. G., T. Z., and P. M.; Investigation: A. G., T. Z., C. G., P. M., H. C., K. S., C. P., E. B., and T. H. N. ; Writing-Original Draft: J.d.l.C.Z, A. G., and L. M; Writing-Review & Editing: all, Supervision: K. C. H., and L. M.; Funding Acquisition: L. M.

## Resource Availability

### Lead contact

Further information and requests for resources and reagents should be directed to the lead contact, Lisa Maier.

### Materials availability

Plasmids generated in this study are available from the lead contact upon request.

### Data availability

The 16S rRNA gene amplicon sequencing data generated during this study have been deposited in the European Nucleotide Archive with accession ID PRJEB65315. Any additional information required to reanalyze the data reported in this paper is available from the lead contact upon request.

### Code availability

The R notebooks with the code used for data analysis are available at https://github.com/Lisa-Maier-Lab/HTD_CR.

## Methods

### Bacterial cultivation of monocultures, Com20/21, and stool-derived communities

The species used in this study are listed in the Supplementary Table S10. They were purchased from DSMZ, BEI Resources, ATCC, or Dupont Health & Nutrition, or were generously provided as gifts from the Denamur Laboratory (INSERM), the Blokesch Laboratory (EPFL), the Andrews-Polymenis Laboratory (Texas A&M University), the Darby Laboratory (UCSF), or the Wagner Laboratory (University of Tübingen). All gut commensal species, whether grown individually or as a community, were cultivated in mGAM medium (HyServe GmbH & Co.KG, Germany) at 37 °C, with the exception of *Veillonella parvula* and *Bilophila wadsworthia* monocultures. *V. parvula* was cultured in Todd-Hewitt Broth supplemented with 0.6% sodium lactate and *B. wadsworthia* was cultured in mGAM supplemented with 60 mM sodium formate and 10 mM taurine. The media were pre-reduced for a minimum of 24 h under anoxic conditions (2% H_2_, 12% CO_2_, 86% N_2_) in an anaerobic chamber (Coy Laboratory Products Inc.). The species were inoculated from frozen stocks into liquid culture media and passaged twice (1:100) overnight to ensure robust growth. The purity and identity of the species was regularly verified via sequencing of the 16S rRNA gene and/or MALDI TOF mass spectrometry (MS)^45^.

A representative set of prevalent and abundant species from the human gut microbiome was selected as previously described^10,46^. From this set, 31 phylogenetically diverse species, differing >3 % in their 16S rRNA gene sequences in the V4 region, were selected. When monocultures of all species were mixed in equal ratios, 20 of these species were consistently detectable and their levels were stable after several passages^17^.

For experiments involving human stool-derived material informed consent was obtained from all 8 donors (approved by the Ethics Committee of the University Hospital Tübingen, project ID 314/2022B02). Stool-derived communities were generated from fresh human fecal samples as previously described^29,30^ by inoculating from frozen glycerol stocks into 3 mL of BHI and serial dilution of 1:200 for 3 48-h passages to ensure that composition reached a steady state before measurements. Experiments were performed in clear, flat-bottomed 96-well plates (Greiner Bio-One). Plates were sealed with breathable AeraSeals (Excel Scientific). Communities were stored with glycerol as frozen stocks at -80 °C, inoculated from frozen stocks into fresh mGAM and grown overnight.

Selective plating of pathogens was performed under aerobic conditions. For animal experiments, *S*. Tm was cultured in LB broth supplemented with 0.3 M NaCl, and *S*. Tm loads in intestinal contents and organs were determined on MacConkey agar supplemented with 50 µg/mL streptomycin.

### Prestwick library screening for pathogens

Prestwick library screening was performed as previously described^10^ on five pathogenic bacterial species in mGAM medium under anaerobic conditions. In brief, the library, which consists of approximately 1200 FDA-approved drugs, was diluted to 100-fold the working concentration in DMSO (2 mM) in V-bottom polypropylene plates (Greiner Bio-One, cat. No. 651261). For the screening experiments, drug master plates were diluted to 2-fold the working concentration in mGAM (40 µM) in U-bottom plates (ThermoScientific, cat. No. 168136), aliquoted (50 µL per plate) and stored at - 20 °C for a maximum of one month. The DMSO control wells within each 96-well plate served as controls. For experiments with *H. parainfluenzae*, mGAM was supplemented with 0.5 mg/L hemin and 2 mg/L NAD. Prior to inoculation, the drug plates were pre-reduced overnight in an anaerobic chamber.

Before the screening experiments, bacterial strains were passaged twice overnight (1:100) anaerobically and OD at 578 nm (OD_578_) was adjusted to 0.02. After inoculation, the starting OD_578_ for all bacterial species was 0.01 and the drug concentration in the plate was 20 µM with 1% DMSO. All plates were sealed with breathable membranes (Breathe-Easy, Sigma-Aldrich, cat. No. Z380059). Bacterial growth was tracked by measuring the OD_578_ every hour for 24 h using a microplate spectrophotometer (EON, Biotek) coupled with a Biostack 4 microplate stacker (Biotek), both housed inside an incubator (EMBL workshop). All screening experiments were performed in three biological replicates. For analysis, growth curves were truncated at the transition from exponential to stationary phase for analysis. The area under the curve (AUC) was calculated using the trapezoidal rule and normalized to the solvent/DMSO controls within the same plate. We identified hits from normalized AUC measurements by fitting heavy-tailed distributions, specifically the scaled Student’s t-distribution^47^, to the wells containing controls. P values for each drug and strain were combined across replicates using Fisher’s method, and the False Discovery Rate (FDR) was calculated using the Benjamini-Hochberg method over the entire matrix.

### Drug selection

Drugs for the *in vitro* challenge assay were selected based on their direct inhibitory effect on members of Com20^10^. We aimed to identify drugs with different inhibition profiles across the 20 species so that we could generate communities with sufficient compositional variation. We performed hierarchical clustering (Euclidean distance metric and complete linkage method) using the normalized AUC values of the 172 drugs that showed significant inhibition (adjusted P value < 0.01) against at least 5 of the 20 species in Com20. From these clusters, we selected 30 drugs representing diverse inhibition spectra across Com20 members. Additionally, we incorporated 25 clinically relevant drugs, resulting in a final selection of 52 drugs. None of the drugs interfered with the luminescence readout of the assay. Drugs directly inhibited *S*. Tm growth were excluded. We further excluded beta-lactam antibiotics due to the presence of ampicillin resistance on the pilux plasmid used in the *S*. Tm invasion assay. Among this final set, there were 43 human-targeted drugs from distinct therapeutic classes and 7 antibiotics (Supplemantary Table S3). For each drug, we tested five concentrations. Drugs with reported intestinal concentrations exceeding 20 µM^10^ were screened at concentrations from 10 to 160 µM, and the remaining drugs were screened at concentrations from 2.5 to 40 µM.

### IC_25_ determination

All drugs were dissolved in DMSO, except for clomipramin, doxorubicin, and tobramycin, which were dissolved in water. Drug master plates at a concentration 100 times the working concentration were prepared by serially diluting the stock solutions two-fold in DMSO or water. The dilutions were carried out column-wise in V-bottom 96-well plates (Greiner Bio-One, cat. No. 651261), starting from 160 mM. Each column in the plate contained 8 two-fold dilutions of a drug, except for column 7 containing DMSO or water as a control. This strategy resulted in 11 drugs screened per plate. The master plates were diluted to 2 times the assay concentration in 50 µL of mGAM in U-bottom 96-well plates (Thermo Fisher Scientific, cat. No. Z168136) and stored at -20 °C for a maximum of one month. Prior to the assay, the plates were thawed and pre-reduced overnight in an anaerobic chamber.

Monocultures or stool-derived communities^24^ were grown overnight in 5 mL of mGAM. The next day, they were diluted to OD_578_ of 0.02. Then, 50 µL of this suspension were added to the drug plates to result in a starting OD_578_ of 0.01 and a DMSO concentration of 1% in all wells. Plates were sealed with a Breathe-Easy breathable membrane (Sigma-Aldrich, cat. No. Z380059). Growth curves of OD_578_ were monitored every hour after 1 min of linear shaking under anaerobic conditions using an Epoch2 microplate reader coupled with a Biostack 4 microplate stacker (both Agilent) housed in a custom-made incubator (EMBL workshop^24^). At least three biological replicates were analyzed for each species.

To calculate the AUC, the growth curves were analyzed as previously described^10,24^ with the R package ‘neckaR’ (https://github.com/Lisa-Maier-Lab/neckaR), using control wells within the plate that did not contain any drugs to define normal growth. The median AUC was calculated for each concentration across the three replicates. To conservatively remove the effects of noise, monotonicity was enforced. If the AUC decreased at lower concentrations, it was set to the highest AUC measured at higher concentrations. The IC_25_ was defined as the lowest concentration at which a median AUC <0.75 was observed.

### *In vitro* invasion assay for *S.* Tm

#### Preparation of drug master plates

Drug master plates were prepared in V-bottom 96-well plates at 100-fold the drug working concentration in DMSO as described above. Concentration gradients of the drugs (16 mM-1 mM or 4 mM-0.25 mM) were represented by each column, with row B and G having the highest and lowest concentration, respectively. In each deep-well plate, Row E each served as solvent controls containing only DMSO or water. To prepare the 96-deep-well plates (Thermo Fisher Scientific, cat. No. AB-0564) for the *S*. Tm challenge assay, 5 µL of the drug master plate were transferred to the deep-well plate, which already contained 95 µL of mGAM medium. Subsequently, the deep-well plates (5 times the drug working concentration in 5% DMSO) were pre-reduced overnight in an anaerobic chamber. Wells on the border contained only mGAM media (sterile controls).

#### Assembly of Com20/Com21

For Com20/Com21 assembly, each member was inoculated from frozen stocks and cultured anaerobically in 5 mL of mGAM over two overnight passages (1:100) as monocultures. OD_578_ was individually measured for each species. The cultures were mixed together in the volume required to achieve a total OD_578_ of 0.0125 (e.g., in Com20, each species contributed an OD_578_ of 0.000625) and 400 µL of this suspension were added to wells of 96-deep-well plates containing drugs as described above to achieve a starting OD_578_ of 0.01 (total volume of 500 µL).

The deep-well plate with the drugs and communities was sealed with an AeraSeal breathable membrane (Sigma-Aldrich, cat. No. A9224) and incubated anaerobically at 37 °C for 24 h. The 24 h of incubation with drugs disrupted the composition of the communities, which were used for the *in vitro S*. Tm challenge assay. Pellets from 300 µL of the cultures were frozen for 16S rRNA gene analysis.

#### In vitro S. Tm challenge assay

For the luminescence-based invasion assay, we used the human gut pathogen *Salmonella enterica* serovar Typhimurium strain SB300^48^ with the plasmid pIJ11282 ilux (pRS16591, *S*. Tm pilux; gift from Foster Lab, University of Oxford) for constitutive expression of the ilux operon under the nptII promoter^49^. *S*. Tm pilux was grown anaerobically at 37 °C overnight in mGAM supplemented with 100 µg/mL ampicillin and then sub-cultured by diluting 1:100 in the same medium. The next day, we measured the OD_578_ of 100 µL of all drug-perturbed communities in a 96-well clear, flat-bottom plate. To assess the growth potential of *S*. Tm pilux in the drug-perturbed communities, 50 µL from each well of the drug-perturbed communities were transferred into new pre-reduced, deep-well plates. *S*. Tm pilux was diluted to an OD_578_ of 0.0025 and 200 µL of this suspension were added to the assay deep-well plate. Two hundred fifty microliters of mGAM were added so that the total volume was 500 µL, resulting in a starting OD_578_ for *S*. Tm of 0.001 and for the untreated community of 0.5. The assay plate was sealed with an AeraSeal breathable membrane (Sigma-Aldrich, cat. No. A9224) and incubated anaerobically at 37 °C for 4.5 h. Thereafter, the plate was taken out of the anaerobic chamber. The contents of the wells were thoroughly mixed and 25 µL of 2 mg/mL chloramphenicol were added to each well to halt *S*. Tm growth and stabilize the luminescence signal. One hundred microliters of the cell suspension were transferred to a white 96-well plate (Thermofisher 236105). Approximately 10 min later, the plate was incubated for 10 min at 37°C in a Tecan Infinite 200 PRO microplate reader and luminescence was measured.

The *in vitro S.* Tm challenge assay involved testing 260 conditions, consisting of 52 drugs at 5 different concentrations, and was performed in triplicates. We obtained two measurements: the OD_578_ of communities after overnight incubation with the drugs, and the luminescence emitted by *S*. Tm as a proxy for pathogen growth within the drug-perturbed communities. For data analysis, OD_578_ values were first corrected by subtracting the baseline OD_578_ from mGAM medium. Subsequently, the luminescence and OD_578_ values were normalized to the control column in row E, which contained the unperturbed community (solvent controls). Both *S*. Tm luminescence and Com20 OD_578_ were highly correlated among the three replicates (R^2^=0.63-0.76 and R^2^=0.86-0.92, respectively; Supplemantary Figure 2e).

#### Variations of the in vitro S. Tm challenge assay: pairwise co-culture and single-species dropout assays

Pairwise co-culture assays were conducted to measure the contribution of each member of Com20 to *S*. Tm growth alone. The commensal strain and *S*. Tm pilux were grown anaerobically overnight in mGAM and subcultured once before the experiment. On the following day, the commensal and *S*. Tm pilux were mixed in 96-deep-well plates with a total volume of 500 µL of mGAM medium. The initial OD_578_ of the commensals was set to 0.1, while *S.* Tm had an initial OD_578_ of 0.0002 (commensal to pathogen ratio 500:1), as described above for the *S*. Tm challenge assay in Com20. Control wells contained only *S*. Tm in monoculture. After a growth period of 4.5 h at 37 °C under anaerobic conditions, *S*. Tm levels were calculated as described for the *in vitro S*. Tm challenge assay.

The single-species dropout assay was performed similar to the *in vitro* S. Tm challenge assays, but one strain at a time was omitted when assembling the synthetic community.

#### Quantification of S. Tm in treatment-mimicking communities

To validate our screen, we selected four conditions (treatment with erythromycin, floxuridine, sertindole, and zafirlukast). Based on the composition of Com20 after treatment with these drugs, we assembled treatment-mimicking communities containing only the members with a mean relative abundance 23% after 24 h of drug exposure. Com20 and treatment-mimicking communities were grown in a deep-well plate at 37 °C anaerobically. After 24 h, dilution series of these communities were performed in a deep-well plate in a total volume of 400 µL. We transferred 100 µL from each dilution and Com20 to a flat-bottom plate and measured the OD_578_. Fifty microliters were transferred to a new deep-well plate containing 250 µL mGAM per well. In addition, 200 µL of *S*. Tm pilux (OD_578_ 0.0025) were added to each well and the plate was incubated for 4.5 h at 37 °C. The remaining volume of the dilution series was kept for DNA isolation and 16S rRNA gene sequencing. After 4.5 h, *S*. Tm levels were determined via luminescence measurements as described above. The experiment was performed in triplicate and luminescence values detected in treatment-mimicking communities were normalized to luminescence values detected in Com20. We compared the log2-fold change of *S*. Tm luminescence and OD578 between treatment-mimicking communities and drug-treated communities to identify the dilution step that best mimicked the drug-treated community.

#### Analysis of community composition using 16S rRNA gene amplicon sequencing

DNA was extracted from pellets of 300 µL culture using a DNeasy UltraClean 96 Microbial Kit (Qiagen 10196-4) or from whole fecal pellets using a DNeasy PowerSoil HTP 96 kit (Qiagen 12955-4). Library preparation and sequencing was performed at the NGS Competence Center NCCT (Tübingen, Germany). Genomic DNA was quantified with a Qubit dsDNA BR/HS Assay Kit (Thermo Fisher) and adjusted to 100 ng input for library preparation. The first step PCR was performed in 25 µL reactions including KAPA HiFi HotStart ReadyMix (Roche), 515F^50^ and 806R^51^ primers (covering a ∼350-bp fragment of the 16S V4 region) and template DNA (PCR program: 95 °C for 3 min, 28X (98 °C for 20 s, 55 °C for 15 s, 72 °C for 15 s), 72 °C for 5 min). Initial PCR products were purified using 28 µL of AMPure XP beads and eluted in 50 µL of 10 mM Tris-HCl. Indexing was performed in a second step PCR including KAPA HiFi HotStart ReadyMix (Roche), index primer mix (IDT for Illumina DNA/RNA UD Indexes, Tagmentation), purified initial PCR product as template (PCR program: 95 °C for 3 min, 8X (95 °C for 30 s, 55 °C for 30 s, 72 °C for 30 s), 72 °C for 5 min). After another bead purification (20 µL of AMPure XP beads, eluted in 30 µL of 10 mM Tris-HCl), the libraries were checked for correct fragment length on an E-Base device using E-Gel 96 Gels with 2% mSYBR Safe DNA Gel Stain (Fisher Scientific), quantified with a QuantiFluor dsDNA System (Promega), and pooled equimolarly. The pool was sequenced on an Illumina MiSeq device with a v2 sequencing kit (input molarity 10 pM, 20% PhiX spike-in, 2×250 bp read lengths.

### Computational processing of 16S rRNA amplicon sequences

We used the DADA2 v. 1.21.0^52^ package of R (R v. 4.2.0) following its standard operating procedure at https://benjjneb.github.io/dada2/bigdata.html. Briefly, after inspecting the quality profiles of the raw sequences, we trimmed and filtered the paired-end reads using the following parameters: trimLeft: 23, 24; truncLen: 240, 200; maxEE: 2, 2; truncQ: 11. The filtered forward and reverse reads were dereplicated separately and used for inference of amplicon sequence variants (ASVs) using default parameters, after which the reads were merged on a per-sample basis. Next, we filtered the merged reads to retain only those with a length between 250 and 256 bp and carried out chimera removal.

We performed the taxonomic assignment in two steps. First, the final set of ASVs was classified up to genus level using a curated DADA2-formatted database based on the genome taxonomy database (GTDB) release R06-RS202^52^ at https://scilifelab.figshare.com/articles/dataset/SBDI_Sativa_curated_16S_GTDB_database/14869077. Next, ASVs belonging to genera expected to be in COM20 were further classified at the species level using a modified version of the aforementioned database that contained only full length 16S rRNA sequences of the 20 members of the synthetic community. The sequence of each ASV was aligned against this database using the R package DECIPHER v. 2.24.0^53^; we classified an ASV as a given species if it had sequence similarity >98% to the closest member in the database. The abundance of each taxon of COM20 was obtained by aggregating reads at the species level. ASVs from *in vitro* communities and gnotobiotic mice were classified using the two-step processes; ASVs from SPF mice were classified using only the first step. We removed potential contaminant sequences from SPF mouse samples using the permutation filtering method implemented in the PERFect v. 1.14.0 R package^54^; on average, we retained 92.3% of the original sequencing reads (range 77.2%-99.1%).

### Classification of drug treatments according to *S.* Tm growth

We grouped treatments according to their effect on COM20 *in vitro.* To do so, we calculated the mean normalized luminescence and the 95% confidence interval (CI) of each drug-concentration combination. A treatment was classified as *’S.* Tm-favoring’ if its mean normalized luminescence was >2 and the 95% CI did not span 2, the treatment was classified as *’S.* Tm -restricting’ if the mean normalized luminescence was <0.5 and the 95% CI did not span 0.5, and the treatment was classified as having ‘No effect’ if the mean normalized luminescence was between 0.5 and 2. Communities with a normalized OD_578_ <0.2 were also classified but marked for removal in downstream analyses to minimize the bias introduced by low-biomass samples.

### Assessment of drug treatment on the microbiome of synthetic communities *in vitro*

We assessed differences in the composition of the microbial communities among colonization groups (i.e., ‘*S.* Tm-favoring’, ‘*S.* Tm-restricting’, and ‘No effect’). We transformed the ASV abundances using the centered-log-ratio (clr) to account for the compositional nature of the sequencing data. Positive clr values imply that an ASV is more abundant than average; conversely, negative values imply that the ASV is less abundant than average. Low biomass samples were removed. Next, we fitted linear regression models to determine which species were differentially abundant between each colonization group and controls using the MaAsLin2 v. 1.13.0 R package^55^; we included normalized OD_578_ as a covariate to account for the biomass of the community, with the OD_578_ values of control samples set to 1. P values were adjusted using the Benjamini-Hochberg method and a significance threshold of 0.1 was used.

### Prediction of functional potential of synthetic communities and analysis of differentially abundant functions

We used the 16S rRNA gene amplicon data from control and drug-treated Com20 and Com21 *in vitro* communities and untreated gnotobiotic mice samples to predict the metabolic potential of the microbial communities using PICRUSt2^32^ v. 2.4.1. PICRUSt2 maps the nucleotide sequence of each ASV to a database of genomes, which is used to retrieve the functions encoded by each of the detected taxa in a microbial community. Since the composition of the synthetic communities is known, we retrieved the full-length sequences of the 16S rRNA gene for each of the member species and used them together with the species abundance data to predict metagenome functions. For our analyses, we used MetaCyc pathway abundances.

We compared the number of metabolic pathways detected in the untreated *in vitro* and *in vivo* synthetic communities to actual human gut metagenomes. To do so, we retrieved publicly available tables of MetaCyc pathway abundances processed using HUMAnN2 from https://github.com/gavinmdouglas/picrust2_manuscript/tree/master/data/mgs_validation. These tables comprised 156 samples from the Human Microbiome Project^56^ and 57 from Cameroon^57^. For each set of samples, we considered a pathway as present if it was detected in 220% of samples.

For downstream analyses we used the predicted abundance of MetaCyc pathways, which we transformed using the clr to account for the compositional nature of the sequencing data. Positive clr values imply that the pathway is more abundant than average; conversely, negative values imply that the pathway is less abundant than average. Low biomass samples were removed.

We calculated differences in functional beta diversity between colonization groups (i.e., ‘*S.* Tm-favoring’, ‘*S.* Tm-restricting’, and ‘No effect’) and untreated controls with a permutational multivariate analysis of variance (PERMANOVA) test on Aitchison’s distance matrices (Euclidean distance using clr-transformed abundances) using the vegan v. 2.6-4 R package^58^. We performed pairwise PERMANOVA tests contrasting each treatment group to untreated controls accounting for normalized OD_578_ in the models; the OD_578_ value of control samples was set to 1. P values were adjusted using the Benjamini-Hochberg method and a significance threshold of 0.1 was used. We then fitted linear regression models to determine which predicted pathways were differentially abundant between treatment groups compared to controls using the MaAsLin2 v. 1.13.0 R package^55^. We used clr-transformed abundances and included the normalized OD_578_ as a covariate to account for community biomass, the OD_578_ value of control samples was set to 1. P values were adjusted using the Benjamini-Hochberg method; since this analysis is based on a bioinformatics prediction and not an actual metagenome measurement, we used a more stringent significance threshold of 0.01.

### Evaluation of metabolic overlap between *S.* Tm and members of the synthetic community

We estimated potential niche overlap between *S.* Tm and each member of COM21 by calculating the competition and complementarity indices using PhyloMint^26^ v. 0.1.0. The metabolic competition index is a proxy of the metabolic overlap of two species; this index is a non-symmetric measure, and it is calculated based on the number of compounds required but not synthesized by both species^59^ Conversely, the metabolic complementarity index is a proxy for potential syntrophy between species; this index is a non-symmetric measure calculated based on the number of compounds that one species produces that the second species requires but cannot synthesize^59^. Briefly, PhyloMint takes as input the whole genome sequence of each strain, which it uses to obtain a genome-scale metabolic model with CarveMe^25^ v. 1.1, extract the metabolite seed sets, and calculate the competition and complementarity indices.

### *in vitro* challenge assay for other pathogenic *Enterobacteriaceae*

#### Plasmid transformation

Bacterial strains were grown overnight in 6 mL of LB medium at 27 °C (WA-314, YpsIII) or 37 °C (*Kp* MKP103, *Ec* CFT073). Overnight cultures were centrifuged for 5 min at 4000*g*. Pellets were washed twice with 5 mL of 300 mM sucrose solution, transferred in 1 mL of 300 mM sucrose solution to an Eppendorf cap, and centrifuged for 1 min at 10,000*g*. The supernatant was removed, the bacteria were resuspended in 100 µL of 300 mM sucrose solution and transferred to a Gene Pulser Cuvette (0.2-cm electrode gap, Bio-Rad), and 100 ng of plasmid DNA (pEB1GM or pEB2GO) were added. Subsequently, electroporation was performed using a Gene Pulser (Bio-Rad) and 1 mL of LB was immediately added. The bacterial suspension was then shaken at the appropriate temperature for 1 h and plated on LB-gentamicin plates overnight. Successful electroporation was verified by measuring chemiluminescence of the lux reporter.

#### Adaptations of the S. Tm challenge assay to other pathogens

Other *Gammaproteobacteria* species were screened similarly to *S*. Tm in the challenge assay described above. Drug master plates were prepared in the same manner, except that only 12 drugs were tested and the master plate concentration ranged from 10 mM to 1 mM. Moreover, only the outer rows were left empty to serve as medium controls. Post-drug expansion of other *Gammaproteobacteria* was only tested in COM20, which was assembled as described above. For the luminescence-based assay, we used the human gut pathogens *Escherichia coli* CFT073, *Klebsiella pneumoniae* MKP103, *Shigella flexneri* 24570, *Yersinia enterocolitica* WA-314, *Yersinia pseudotuberculosis* YPIII, and *Vibrio cholerae* A1552. Except for *V. cholerae*, all pathogens contained a variant of the pilux plasmid that enabled constitutive expression of the lux reporter. All pathogens were grown anaerobically at 37 °C overnight in mGAM supplemented with 100 µg/mL ampicillin (*S. flexneri*), 15 µg/mL gentamicin (*E. coli*, *Y. enterocolitica*, *Y. pseudotuberculosis*) or 75 µg/mL gentamicin (*K. pneumoniae*) and then sub-cultured by diluting 1:100 in the same medium. We continued as described in the above section on the *in vitro S.* Tm challenge assay but incubated the plates at 37 °C for a species-specific amount of time (4.5 h for *E. coli*, 5 h for *S. flexneri* and *K. pneumoniae*, 5.5 h for *V. cholerae*, 7 h for *Y. enterocolitica* and *Y. pseudotuberculosis*).

For *V. cholerae*, the plates were serially diluted (10^1^-10^8^ fold) in PBS and selectively plated aerobically on LB agar with 100 µg/mL ampicillin for pathogen enumeration. For the other pathogens, their levels were determined as described for *S*. Tm pilux with a Tecan Infinite 200 PRO microplate reader.

The *in vitro* pathogen challenge assay involved testing 60 conditions, consisting of 12 drugs at 5 concentrations, and was performed in triplicate. We obtained two measurements: the OD_578_ of communities after overnight incubation with the drugs and the luminescence emitted by the pathogens (CFU in the case of *V. cholerae*) as a proxy for pathogen growth within the drug-perturbed communities. For data analyses, both the luminescence (CFU for *V. cholerae*) and OD_578_ values were normalized to the median of the controls in row E, which contained the unperturbed community (solvent controls).

### *In vivo* colonization assays for *S*. Tm

Animal experiments were approved by the local authorities in Tübingen (Regierungspräsidium Tübingen, H02/20G and H02/21G). Five to six week-old mice were used and randomly assigned to experimental groups.

#### Defined colonized mice

Germfree C57BL/6J mice were bred in house (Gnotobiotic Mouse Facility, Tübingen), housed under germ-free conditions in flexible film isolators (Zoonlab), and transferred to the Isocage P system (Tecniplast) to perform experiments. We supplied mice with autoclaved drinking water and y-irradiated maintenance chow (Altromin) ad libitum. Female (*n* = 25) and male (*n* = 14) mice were kept in groups of 3-4 animals and were scored daily for their health status.

#### Specific pathogen free mice

Male specific pathogen free (SPF) C57BL/6J mice (cat. no. 632C57BL/6J) were purchased from Charles River Laboratories (Sulzfeld, Germany, Room A004) at the age of 35-41 days. After delivery, mice were kept in groups of 3 in individually ventilated cages (IVC) and had a 2-week acclimatization period. Mice were supplied with autoclaved drinking water and maintenance diet for mice (Sniff) ad libitum. We performed the experiments in a laminar flow system (Tecniplast BS60) and scored animals daily.

#### Preparation of the Com20 bacterial community and colonization of germ-free mice

We prepared Com20 under anaerobic conditions (2% H_2_, 12% CO_2_, rest N_2_) in a chamber (Coy Laboratory Products Inc). Consumables, glassware, and media were pre-reduced at least 2 days before inoculation of bacteria. We grew each strain as a monoculture overnight at 37 °C in 5 mL of their respective growth medium. The next day, bacteria were subcultured 1:100 in 5 mL of fresh medium and incubated for 16 h at 37 °C, except *Eggerthella lenta*, which was grown for 2 days. We measured the OD_578_ and mixed bacteria in equal ratios to a total OD_578_ of 0.5 (OD_578_ of 0.025 for each of the 20 strains) in a final volume of 10 mL. After adding 2.5 mL of 50% glycerol (with a few crystals of palladium black (Sigma-Aldrich)), 200 µL aliquots were prepared in 2-mL glass vials (Supelco, Ref. 29056-U) and frozen at -80 °C. Frozen vials were used within 3 months.

To colonize germ-free mice, cages were transferred to an ISOcage Biosafety Station (IBS; Tecniplast) through a 2% Virkon S disinfectant solution (Lanxess) dipping bath. Glycerol stocks of the frozen Com20 community (one per mouse) were kept on dry ice before being thawed during transfer into the IBS. We used the mixtures directly after thawing with a maximal time of exposure to oxygen of 3 min. We colonized mice by oral gavage (50 µL), and gavaged again after 48 h using the same protocol. The IBS was sterilized with 3% perchloracetic acid (Wofasteril, Kesla Hygiene AG).

To monitor *in vivo* stability of Com20 in gnotobiotic mice, we collected fresh fecal samples from every defined colonized mouse after 2, 6, 28, and 57 days after the second colonization. DNA was extracted using the DNeasy PowerSoil HTP 96 Kit and community composition was analyzed via 16S rRNA amplicon sequencing.

#### In vivo S. Tm challenge

The day before infection, we inoculated an *S*. Tm culture in LB broth supplemented with 0.3 M NaCl using colonies from a plate and grew the culture for 12 h on a rotator (Stuart, SB3, speed 9) at 37 °C. Fifty microliters of *S.* Tm were sub-cultured in 5 mL of LB broth supplemented with 0.3 M NaCl and incubated for 3 h in the same conditions. We washed 1 mL of the subculture twice with 1 mL of ice-cold PBS in a 2 mL Eppendorf tube via centrifugation at 4 °C and 14000*g* for 2 min. The pellet was resuspended in 1 mL of ice-cold PBS and kept on ice until oral administration. Mice were infected with a *S*. Tm load of 5×10^6^ CFU in 50 µL of PBS.

#### S. Tm growth inhibition in defined colonized mice

To determine whether Com20 confers colonization resistance in mice, we colonized germ-free mice with Com20 for 28 days. We then treated them with either 50 µL of 25% DMSO (solvent control, full colonization resistance) or with 50 µL of 25% DMSO supplemented with 20 mg streptomycin (no colonization resistance). For comparison, we treated conventional SPF mice with a complex microbiome in the same manner as the mice colonized with Com20. The next day we infected all groups with 50 µL of 5×10^6^ CFU of *S*. Tm. After 16-20 h, mice were euthanized via CO_2_ and cervical dislocation, dissected, and intestinal contents were collected from the colon. We weighed the fecal samples, diluted them in a buffer (2.5 g of BSA, 2.5 mL of Tergitol, 497.5 mL of PBS) and plated the samples on MacConkey agar containing 50 µg/mL streptomycin. After incubation overnight at 37 °C, we counted colonies of *S*. Tm.

#### Treatment with non-antibiotic drugs and infection with S. Tm

Five non-antibiotic drugs were chosen based on the *S.* Tm challenge assay. Clotrimazole (38 mg/kg), zafirlukast (20 mg/kg), chlorpromazine (3 mg/kg), terfenadine (25 mg/kg), and clomiphene (60 mg/kg) were dissolved in 25% DMSO (DMSO + autoclaved drinking water), aliquoted, and stored in 2-mL glass vials (Supelco, Ref. 29056-U) at -80 °C. For every experiment, drugs were freshly prepared. Defined colonized (28 days post colonization) and SPF mice were orally gavaged daily for 6 days with 50 µL of non-antibiotic drug or 25% DMSO. We collected fresh fecal samples directly before the first treatment (day 0) and after 6 days of treatment (day 6), directly before the infection with *S*. Tm.

Fifteen to twenty h later, mice were euthanized by CO_2_ and cervical dislocation, dissected, and intestinal contents were taken from colon and cecum in pre-weighed 2-mL Eppendorf tubes. After weighing the samples, 500 µL of buffer (2.5 g of BSA, 2.5 mL of Tergitol, 497.5 mL of PBS) and one sterile steel ball (Agrolager, art. No RB-5/G20W) per tube were added. Half a spleen, mesenteric lymph nodes, and half a liver lobe were collected in 2-mL Eppendorf tubes containing 500 µL of buffer and one steel ball. All samples were lysed with a TissueLyser II (Qiagen) for 1 min at 25 Hz. Intestinal contents and organs were plated on MacConkey plates supplemented with 50 µg/mL streptomycin. We incubated the plates at 37 °C aerobically for one night and counted colonies the next day to determine CFU/organ.

### Assessment of drug treatment on the microbiome composition of gnotobiotic and SPF mice

We assessed the effect of drugs on the composition of the gut microbiome of gnotobiotic and SPF mice after 6 days of treatment and compared to untreated controls. For this comparison, we carried out an ANCOVA incorporating the abundance of each ASV at day 0 (pre-treatment), thus estimating the baseline adjusted difference between groups at day 6. ANCOVA models were fitted using a multiple linear regression; robust standard errors were calculated using the sandwich v. 3.0-2 R package^60^, which were then evaluated by a coefficient test as implemented in the lmtest v. 0.9-40 R package^61^. To account for the compositional nature of the sequencing data, we transformed ASV abundances with the centered log-ratio. P values were adjusted using the Benjamini-Hochberg method and a significance threshold of 0.1 was used.

## Supplementary Information

### Supplementary Table Legends

Table S1: Prestwick library screen results of pathogenic *Gammaproteobacteria* species, including *H. parainfluenzae*, *S. enterica* serovar Typhimurium, *S. flexneri*, *Y. pseudotuberculosis*, and *V. cholerae*.

Table S2: IC_25_ values for 67 drugs used to treat 5 pathogens and 19 gut commensal species.

Table S3: Results of the *S*. Tm *in vitro* challenge assay in Com20.

Table S4: Altered pathways in *S*. Tm-favoring conditions.

Table S5: Results of the *S*. Tm *in vitro* challenge assay in Com21.

Table S6: Results of the *in vitro* challenge assay for additional pathogenic *Gammaproteobacteria*.

Table S7: Log2-fold change in the luminescence signal of *S*. Tm in Com20, Com21 and stool-derived communities after drug treatment.

Table S8: Differential abundance of bacterial taxa in Com20-colonized drug-treated mice compared to DMSO controls.

Table S9: Differential abundance of bacterial taxa in drug-treated SPF mice compared to DMSO controls.

Table S10: Key resources used in this study.

**Supplementary Figure 1:**
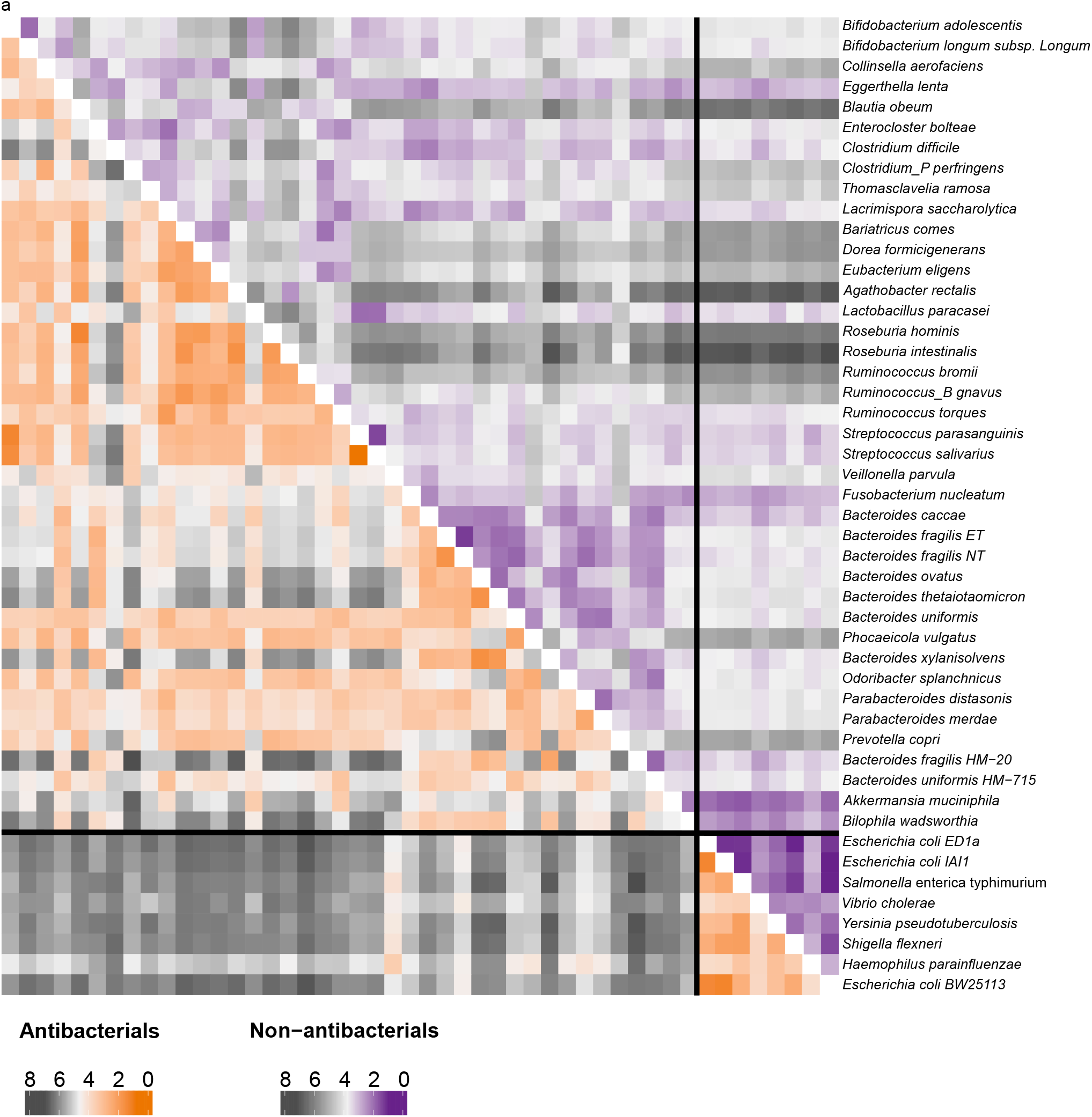
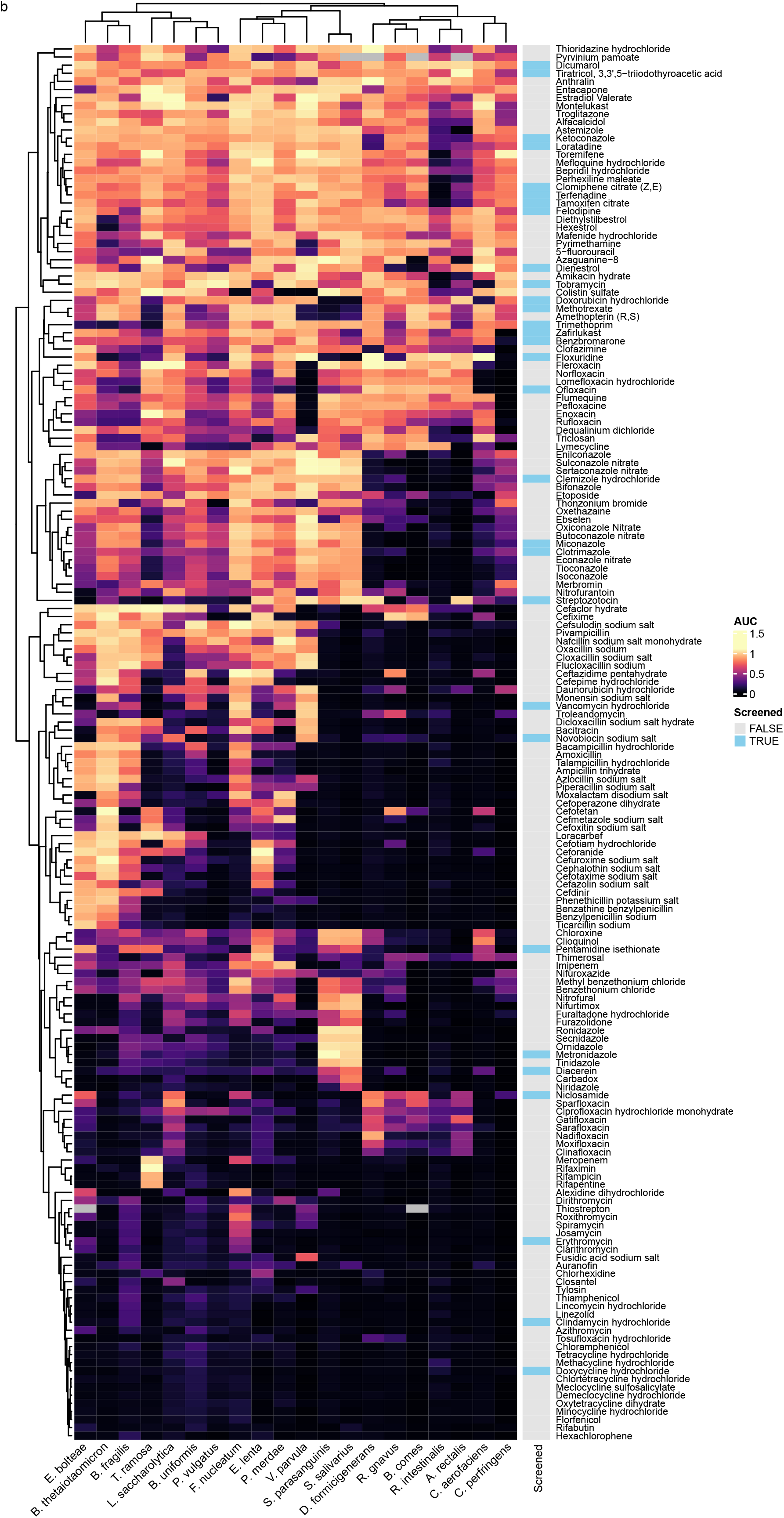
*Gammaproteobacteria* species respond differently to drugs than other gut bacteria. a. Euclidean distances between pairs of bacterial species calculated from the area under the growth curve (AUC) after treatment with 103 antibacterial (lower triangle) and 118 non-antibacterial drugs (upper triangle). On both scales, white corresponds to the median distance across all species, black indicates an overall different response, and orange/violet indicates a similar response across all treatments. Only drugs that inhibited the growth of 25 species were included in the analysis. Species were grouped according to Phylum classification. Note that members of *Gammaproteobacteria*, in the lower right corner, display an overall response similar to each other but distinct to other taxa. b. Heatmap of the AUC for each member of the Com20 community after treatment with 170 drugs, normalized to untreated controls. Color strip indicates whether a drug was included (TRUE, in blue) in our follow-up *S*. Tm challenge assay in Com20. Columns and rows were clustered hierarchically with complete linkage.

**Supplementary Figure 2:**
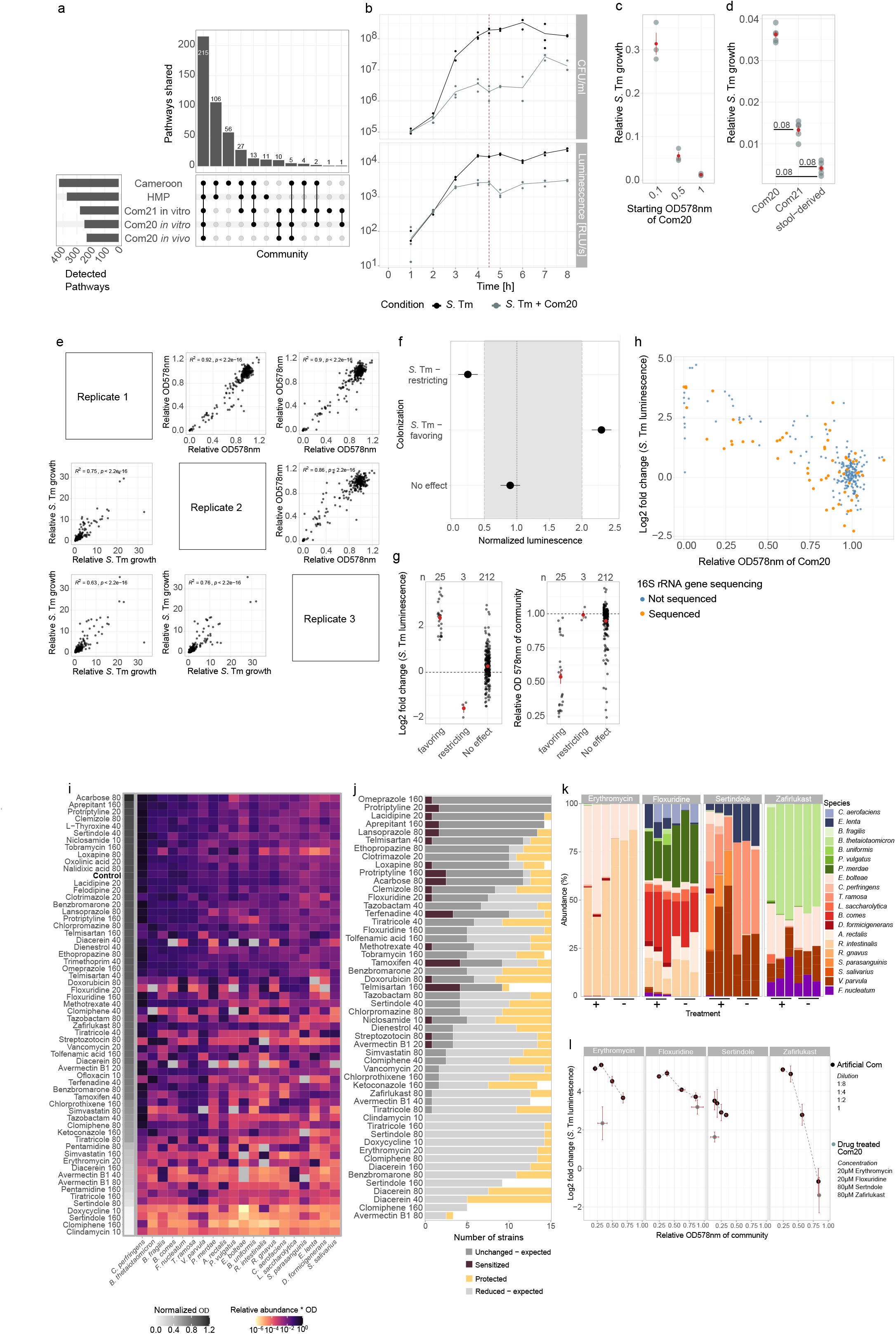
Experimental setup and validation of the *in vitro S.* Tm challenge assay. **a**. Upset plot showing the overlap in the number of MetaCyc metabolic pathways predicted to be present in Com20 (*in vivo* and *in vitro*), Com21 (*in vitro*), and human gut metagenomes from the Human Microbiome Project (HMP) and a Cameroonian cohort. **b***. S*. Tm growth curves based on selective plating (top) or *S*. Tm-specific luminescence (bottom), either alone or within Com20; the line indicates the mean of three biological replicates. At 4.5 h (red vertical line), *S.* Tm growth curves transition to stationary phase. At this time point, luminescence can be used as a proxy of *S.* Tm levels. **c**. Relative growth of *S.* Tm in Com20 compared to pure culture, dependent on the starting OD_578_ of Com20. *S*. Tm was quantified by luminescence. Red points and bars represent the mean of three biological replicates ± SE. **d.** Relative growth of *S.* Tm in Com20, Com21 (consisting of Com20 + *E. coli* ED1a), and a community derived from a fecal sample of a healthy human donor, compared to pure culture. *S*. Tm was quantified by luminescence. Red points and bars represent the mean of three biological replicates ± SE. Adjusted P values from two-sided Wilcoxon-test with correction are shown. **e.** Association of community OD_578_ and *S*. Tm luminescence relative to an untreated community in each of three replicates. *R*^2^ and P values from linear regression models are shown. **f**. Illustrative example of the classification of drug treatments based on relative *S*. Tm growth in the challenge assay; dashed line represents *S*. Tm growth on an untreated community. Points show the mean luminescence of 3 replicates, bars represent the 95 % confidence interval (95% CI). We considered a treatment as *S.* Tm-favoring if its mean normalized luminescence was > 2 and the 95% CI did not span 2; if the mean normalized luminescence was < 0.5 and the 95% CI did not span 0.5, the treatment was classified as *S.* Tm-restricting; if the mean normalized luminescence was between 0.5 and 2 (gray band), the treatment was classified as having ‘No effect’. **g.** Dot plot of normalized *S.* Tm luminescence (left) and community OD_578_ (right) in the challenge assay by treatment group. Black points represent the mean across three replicates. Sample sizes are shown in the top border. Red points and bars represent mean ± SE. **h.** Scatter plot similar to Figure 2c, highlighting samples that were selected for 16S rRNA gene sequencing (57 out of 260) across the distribution of relative community OD_578_ and *S*. Tm luminescence values. **i**. The biomass-scaled relative abundance of each member of a given drug-treated community, calculated by multiplying the normalized OD_578_ of the community by the relative abundance of each taxon from 16S rRNA gene sequencing. The gray-scale column on the left shows the mean OD_578_ of three replicates of the community normalized to an untreated control. **j.** The number of bacterial species displaying emergent (protection or sensitization in community) or expected responses to drug treatments across 52 drugs. An expected response in a community (gray shades) refers to a similar growth pattern (i.e., reduction in growth or similar growth) in monoculture as measured by AUC compared to the OD-scaled relative abundance (i.e., relative abundance * OD) of the microbe as measured by 16S amplicon sequencing; in both cases, measures are normalized to the value of an untreated control. Community protection (yellow) means that the species is affected by drug treatment in monoculture but remains unaltered in the treated community; conversely, community sensitization (burgundy) means that species growth is not affected in monoculture but its abundance decreases in the treated community. Note that the total number of tested species differs between treatments; at the tested concentrations, the analysis of monoculture assays was hampered by erratic bacterial growth patterns, which led to the removal of several growth curves during quality control^24^. **k**. The relative abundances of drug-treated (+) versus drug-mimicking (-) communities for four conditions as determined by 16S rRNA gene sequencing. Each bar represents one biological replicate. **l***. S*. Tm challenge assay in communities mimicking the composition of Com20 after treatment with erythromycin, floxuridine, sertindole or zafirlukast. The communities were diluted to mimic both the composition and the density of the communities after treatment. Dots represent the mean of three replicates, and red lines represent ± SE. Gray dots represent results of the *S*. Tm challenge assay in drug-treated Com20 (replotted from Supplementary Figure 2i) for comparison.

**Supplementary Figure 3:**
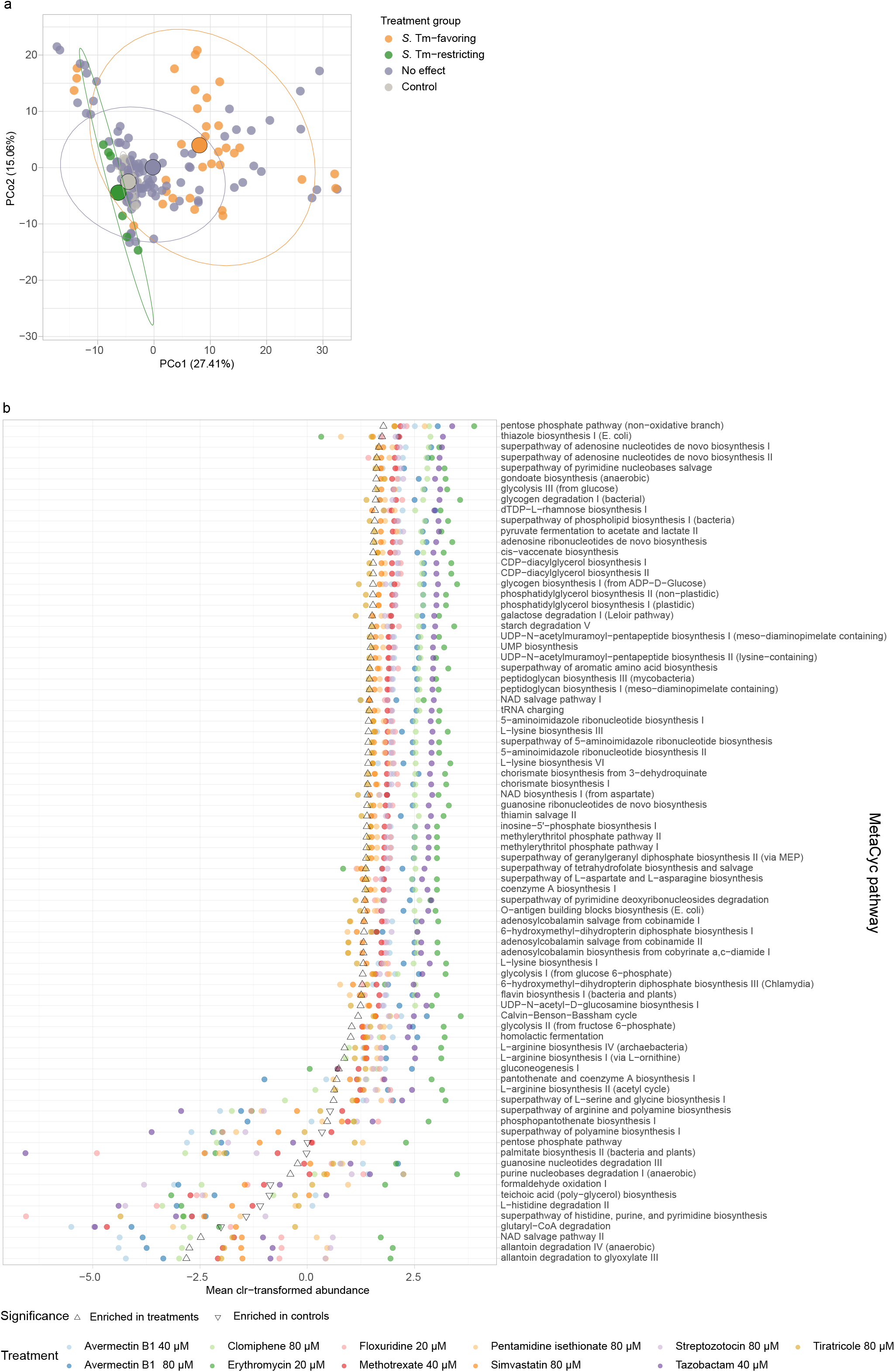
Com20 and *S*. Tm metabolic responses post treatment. **a.** Principal coordinate analysis (PCoA) of drug-treated microbial communities based on Aitchison’s distances calculated from predicted MetaCyc pathway abundances. Colors indicate the *S*. Tm treatment group. Large solid points represent the centroid of each group, ellipses represent 95% confidence intervals. **b.** Dot plot of the mean clr-transformed abundance of predicted MetaCyc pathways that differed significantly between *S*. Tm-favoring treatments compared to untreated controls. Point color indicates treatment, triangles represent the mean abundance in controls, and triangle orientation indicates whether a pathway had a higher abundance in treatments (upward) or in controls (downward).

**Supplementary Figure 4:**
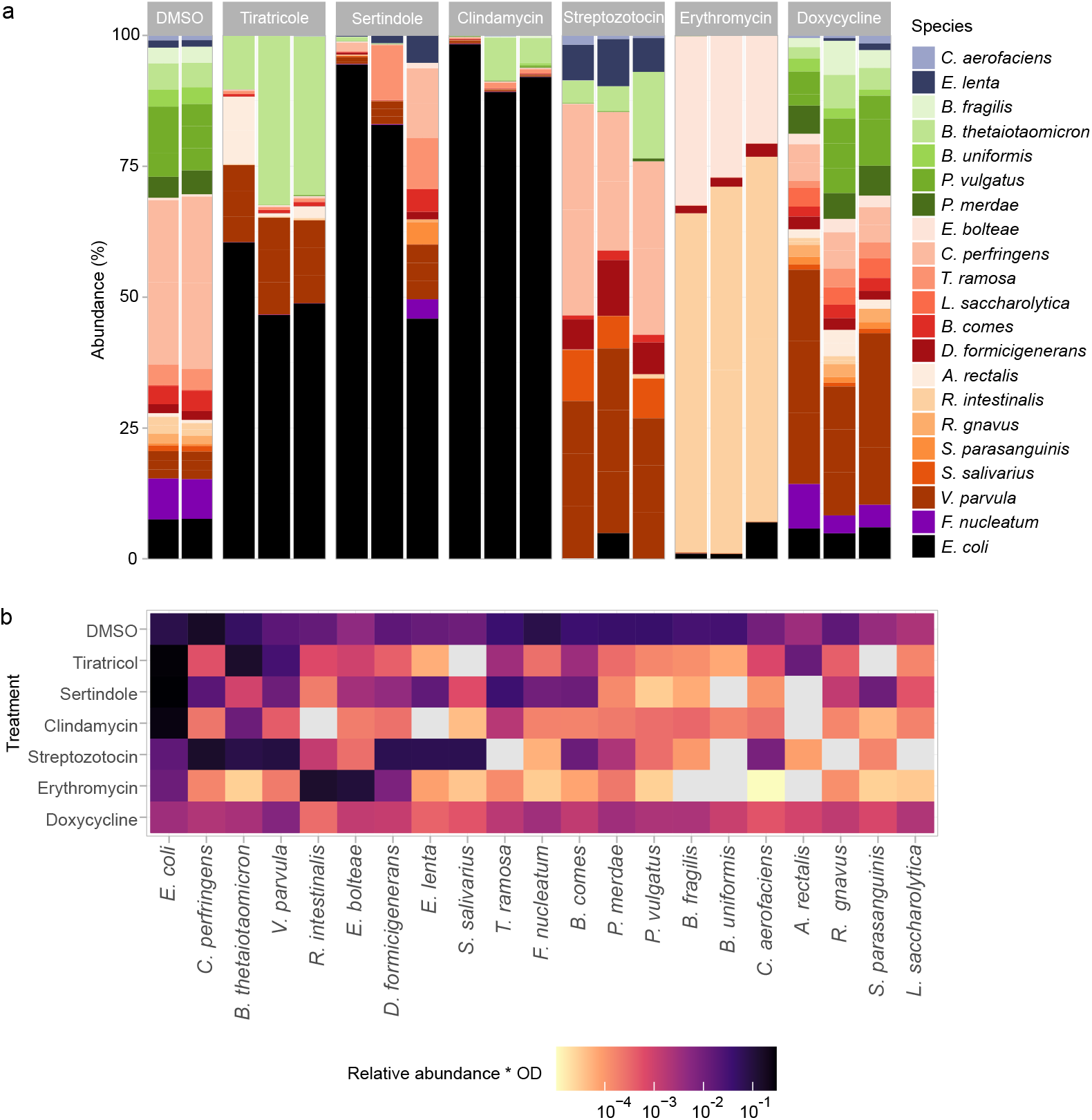
Drug-induced shifts in the composition of Com21. **a.** Relative abundances of taxa in drug-treated Com21 and the DMSO control as determined via 16S rRNA gene sequencing. Each bar represents one biological replicate. **b.** Biomass-scaled relative abundance of each member of drug-treated Com21, calculated by multiplying the normalized OD_578_ of the community with the relative abundance of each taxon obtained by 16S rRNA gene sequencing. Values indicate the mean of three biological replicates.

**Supplementary Figure 5:**
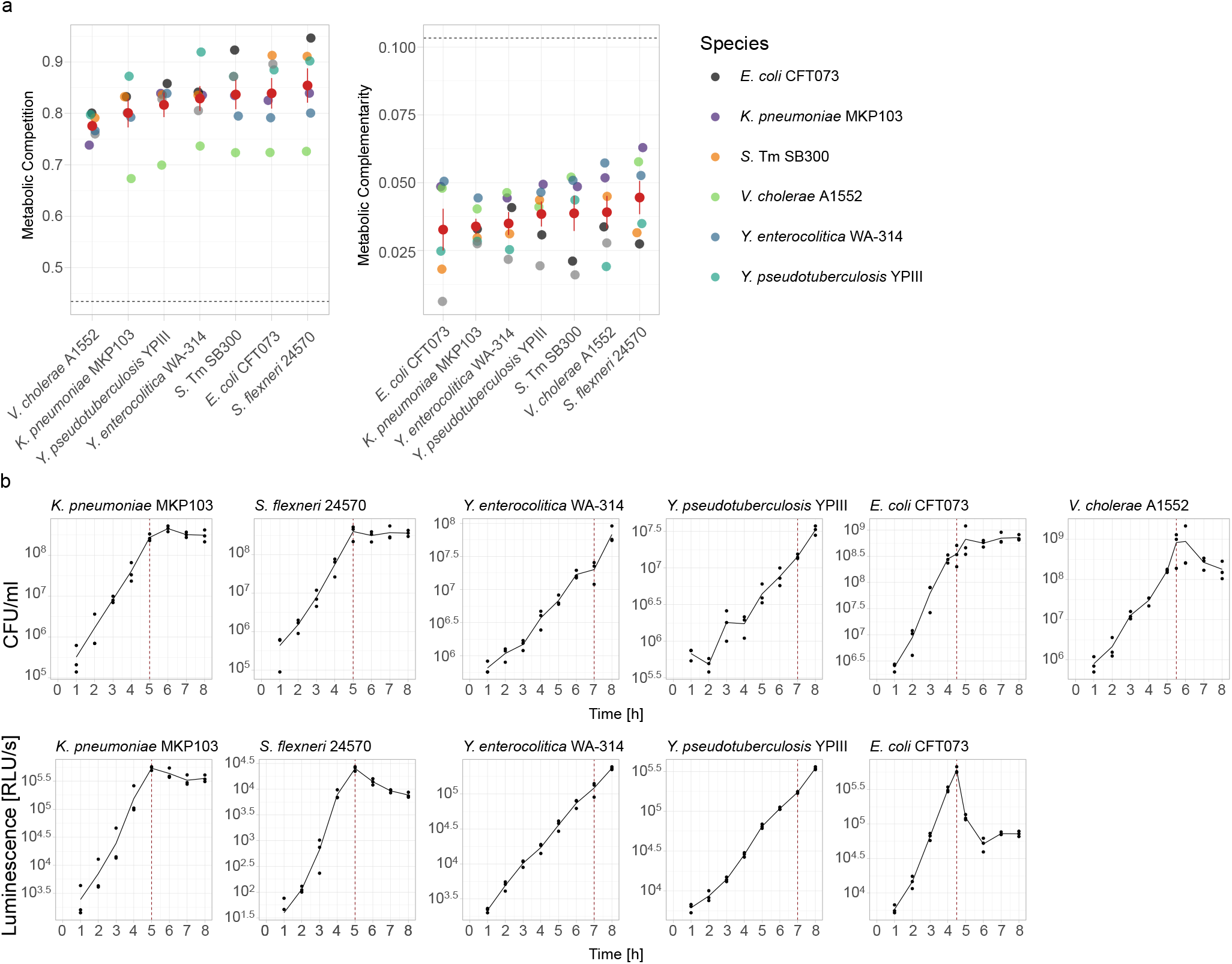
*In vitro* challenge assays for pathogenic *Gammaproteobacteria* species other than *S*. Tm. **a.** Metabolic competition and complementarity indices of pathogenic *Gammaproteobacteria* species, calculated from genome-scale metabolic models. Dashed horizontal lines indicate the mean levels of *S.* Tm with members of Com21, as shown in Figure 4a. Red points and bars represent mean ± SE. Note that the indices are not symmetric. **b**. Growth curves for pathogenic members of *Gammaproteobacteria* based on plating (top) or pathogen-specific luminescence (bottom); the line indicates the mean of 3 biological replicates. Vertical red lines represent the end points selected for pathogen challenge assays. At all endpoints, luminescence can be used as indicators for pathogen levels. *V. cholerae* was enumerated by plating.

**Supplementary Figure 6:**
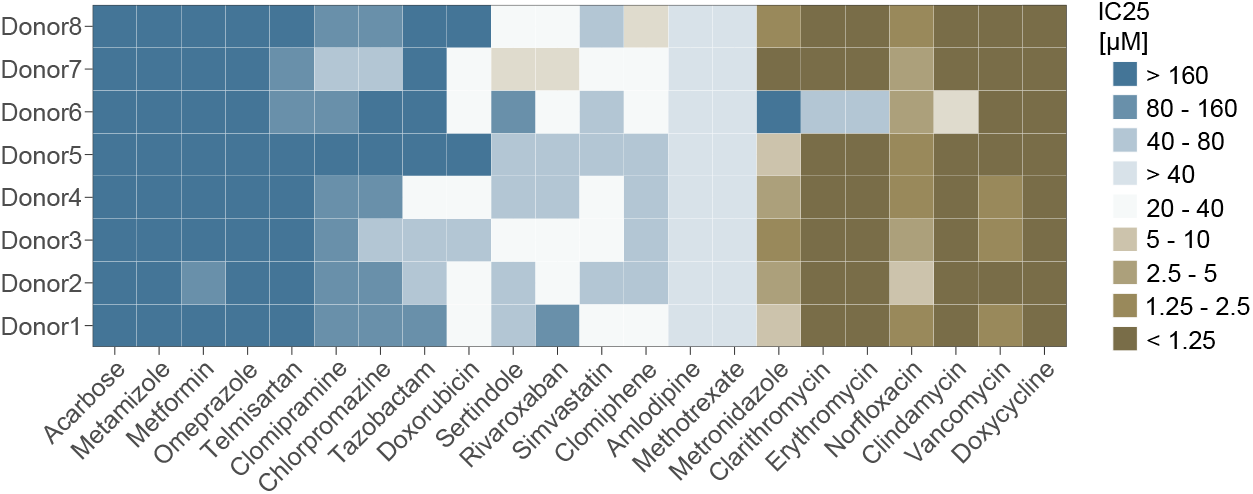
Variation in drug sensitivity between donors. IC_25_ values for 22 drugs used to treat *in vitro* communities derived from the stool of s healthy human donors. Tiles show the mean of 3 biological replicates.

**Supplementary Figure 7:**
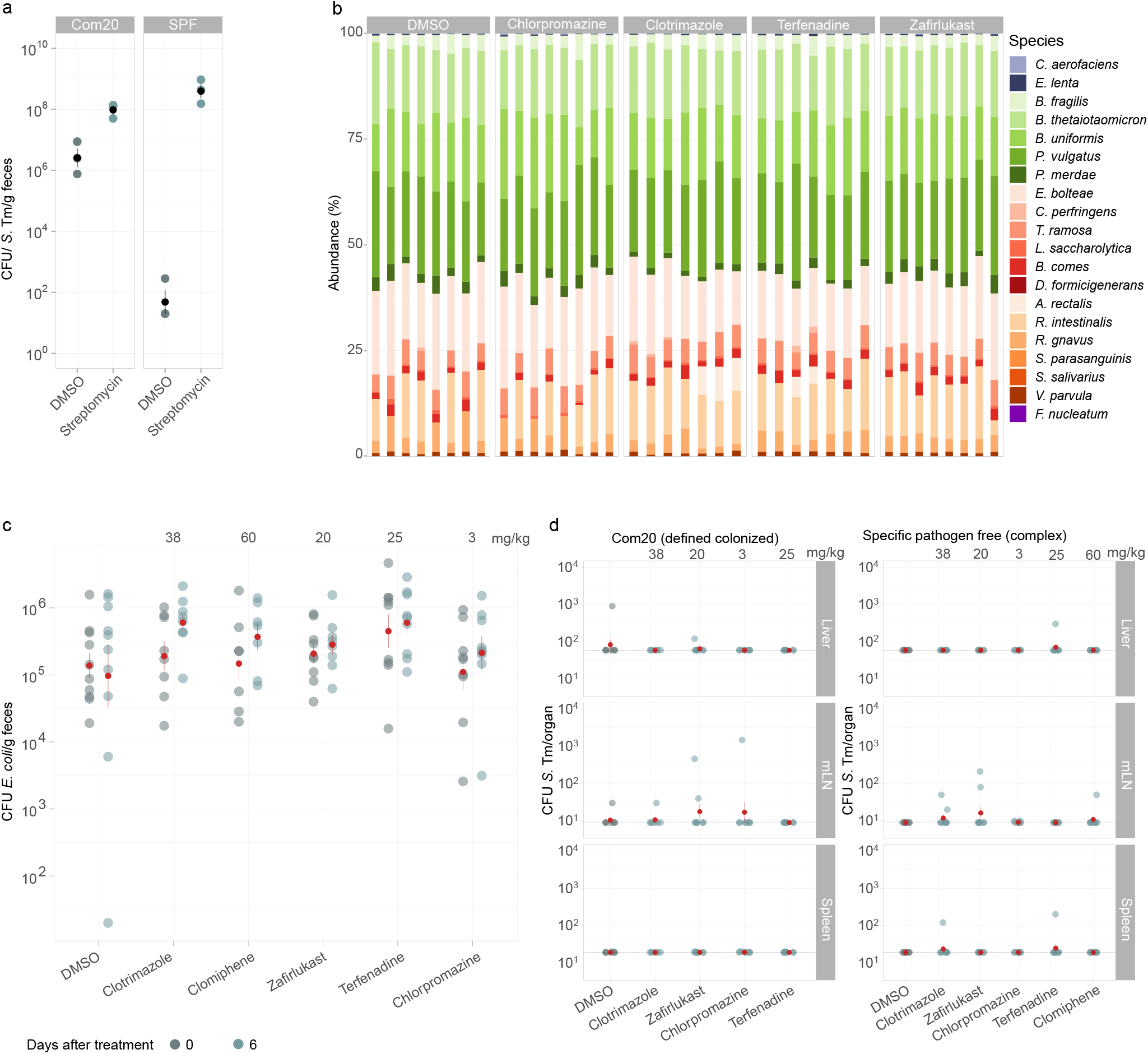
Disruption of colonization resistance by non-antibiotic drugs does not result in systematic spread of *S.* Tm at day 1 post infection. **a**. Fecal *S*. Tm loads on day 1 post challenge with 5x10^6^ CFU *S*. Tm in mice colonized with Com20 compared to conventional SPF mice. Animals were pre-treated either with the vehicle (DMSO) or a single dose of 800 mg/kg streptomycin. Red points and bars represent mean ± SE. **b**. Relative abundances of taxa in feces from Com20-colonized mice after 6 days of treatment. Each column represents the composition of one fecal sample from one mouse. **c***. E. coli* levels in feces of SPF mice before and 6 days after drug treatment as measured by selective plating on MacConkey agar. Drug dosing is indicated on top. Red points show the mean of 7-10 biological replicates and bars represent mean ± SE. Adjusted P values from Two-sided Wilcoxon-test > 0.1 in all cases. **d***. S*. Tm load on day 1 post challenge in the spleen, liver, and mesenteric lymph nodes (mLN). Black horizontal lines indicate detection limits. Red points show the mean of 7-9 biological replicates and bars represent mean ± SE. One-tailed wilcoxon-test with BH correction was performed.

